# Switching perspective: Comparing ground-level and bird’s-eye views for bumblebees navigating dense environments

**DOI:** 10.1101/2023.12.21.572344

**Authors:** Annkathrin Sonntag, Odile Sauzet, Mathieu Lihoreau, Martin Egelhaaf, Olivier Bertrand

## Abstract

Animals navigating in three dimensions encounter different perspectives of their world, often transitioning from bird’s eye views at higher altitudes to ground views closer to the ground. How they integrate this information to pinpoint a goal location is virtually unknown. Here we tested the ability of bumblebees to use both types of views when homing in a dense environment in the vicinity of their inconspicuous nest entrance. Our combined modelling and experimental approach examined various views for localising a goal in dense settings. Whereas, bird’s-eye views performed best in simulations of current nest-centered snapshot homing models, behavioural experiments revealed that bumblebees predominantly relied on ground views when pinpointing nest entrances in dense environments. These findings reveal the limitations of snapshot-homing models and suggest that bumblebees use a combination of navigational tools to successfully find their way home in dense environments. This is not only relevant for understanding bee navigation, but also for other animals and humans navigating in 3D as well as the development of technologies inspired by natural systems, such as autonomous flying robots.

## Introduction

Numerous animals, spanning diverse taxa, navigate within a three-dimensional world where changes in altitude are common. Birds, fish, mammals and insects change their flight altitude, climbing height, or swimming depth from near the ground to great heights above or depths below the water surface during foraging, nesting, or in search for shelter [4, 18, 21, 24, 23, 17]. Also humans experience that the landscapes and views of the environment change with increasing altitude during activities like hiking, climbing, or aeroplane travel. This prompted us to investigate whether animals that have evolved to solve three-dimensional challenges are also adapted to efficiently use bird’s-eye perspectives for navigational purposes. Over the past decades, researchers have delved into the navigational strategies employed by insects, commonly referred to as their navigational toolkit [48]. This toolkit primarily comprises a compass and an odometer, which can be synergistically employed as a path integrator—enabling the integration of distance and direction travelled. Additionally, insects utilise landmark guidance and exploration behaviour in their navigation. Taken all together, the navigational toolkit has been studied by analysing the insects’ walking or flight paths mostly in two dimensions (e.g. [5, 45]).

Accordingly, the visual mechanisms that have been considered are primarily based on the information that can be seen from close to the ground (ground view) [52]. However, when flying insects exit their nest for the first time and are not yet familiar with their surroundings, they increase the distance to the nest during loops, arcs and spirals and their flight altitude during so-called learning flights [11, 30, 36, 39, 49]. They may therefore learn and use visual sceneries at different altitudes. In particular, visual scenery may drastically change when insects change their flight altitude in dense environments with uniformly distributed objects such as grasslands, flower meadows, or forests with similar plants. Flying insects use views from above the dense environment, i.e., bird’ s-eye views, to recognize ground-level landmarks and locations for large-scale navigation [3, 10, 12, 31]. Such bird’s-eye views might not only be relevant at high altitudes and navigation on a large spatial scale, but also on smaller spatial scales and altitudes in the vicinity of the often inconspicuous nest hole during local homing. While several studies have focused on large spatial scales using radar tracking (e.g. [6, 36, 49]), there is limited understanding of the mechanisms behind local homing on a small scale, especially in dense environments.

Using information obtained at different heights might be especially helpful for underground nesting species, such as bumblebees, whose nest entrance is often inconspicuously located within the undergrowth. In such dense environments, bumblebees need to change their flight altitude both when learning the surroundings and when homing. Bird’s eye views might be helpful for guiding homing behaviour in the near-range of the hidden nest hole by providing a kind of overview. In contrast, ground views might help pinpointing the nest entrance. Computational models suggest such views of the visual scenery from within dense environments is sufficient to explain the returning journey of ants [51, 50, 2, 32] and bees [14, 15, 52] under a variety of experimental conditions. These models rely on visual memories of the environments acquired during previous journeys or learning flights at the nest site. To navigate visually, the modelled insect compares its current view with the memorised ones and steers toward the most familiar memories (e.g. active scanning [2, 1], rotation-invariant comparison [43], while oscillating by left and right turns [25, 26]). This class of models can simulate route-following behaviour within dense environments, even at different flight altitudes [37, 8]. Despite all these results at the model level, how insects use views at different altitudes to navigate has never been studied experimentally. Because this is a particular challenge for bumblebees in finding their nest hole, we investigated this in a combined experimental and model analysis.

We addressed the question of whether bumblebees, *Bombus terrestris*, learn views at different altitudes and if they can return home by just using either ground or bird’s eye views. To obtain predictions for the experimental results, we first investigated the navigation of modelled bees guided by standard models (multi-snapshot model), which are frequently used in the literature on homing [53]. We then performed behavioural experiments to challenge bees in the same environment. The analysis was based on homing experiments in object-dense laboratory environments, where the flight altitude during the learning and homing flights was systematically constrained by manipulating the environment. We related our results to the predictions of snapshot models [54, 52] comparing the performance of bird’s and ground view snapshots in the same dense environment like the behavioural experiments.

## Results

### Snapshot models perform best with bird’s eye views

To guide our hypotheses about which views bees should prioritise during their homing flight, we compared predictions of homing models based on either bird’s eye views or ground views. In previous studies on the mechanism of visual homing, models could replicate the insects’ homing behaviour at least in spatially relatively simple situations [7, 2, 26]. These models assume that insects learn one or multiple views, so-called *snapshots*, at or around their goal location, like their nest. During their return, they compare their current views with the stored snapshot(s). Therefore we tested the homing performance in a dense environment of rotational image difference models based on two parameters: brightness values of the pixels [54, 13] or contrast-weighed-nearness values encoding the depth and contrast of the environment based on optic flow information [14] (for details see Methods). The environment for the model simulations was the same as that used in the experimental analysis (Fig. 2 A&B). It consisted of an inconspicuous goal on the floor, i.e. the nest hole, surrounded by multiple, similarly-looking objects creating a dense, artificial meadow around the nest. Thus, we investigated the model performance of homing in a dense environment and carried out an image comparison of the snapshots taken either above the dense environment or within it.

**Figure 1.**
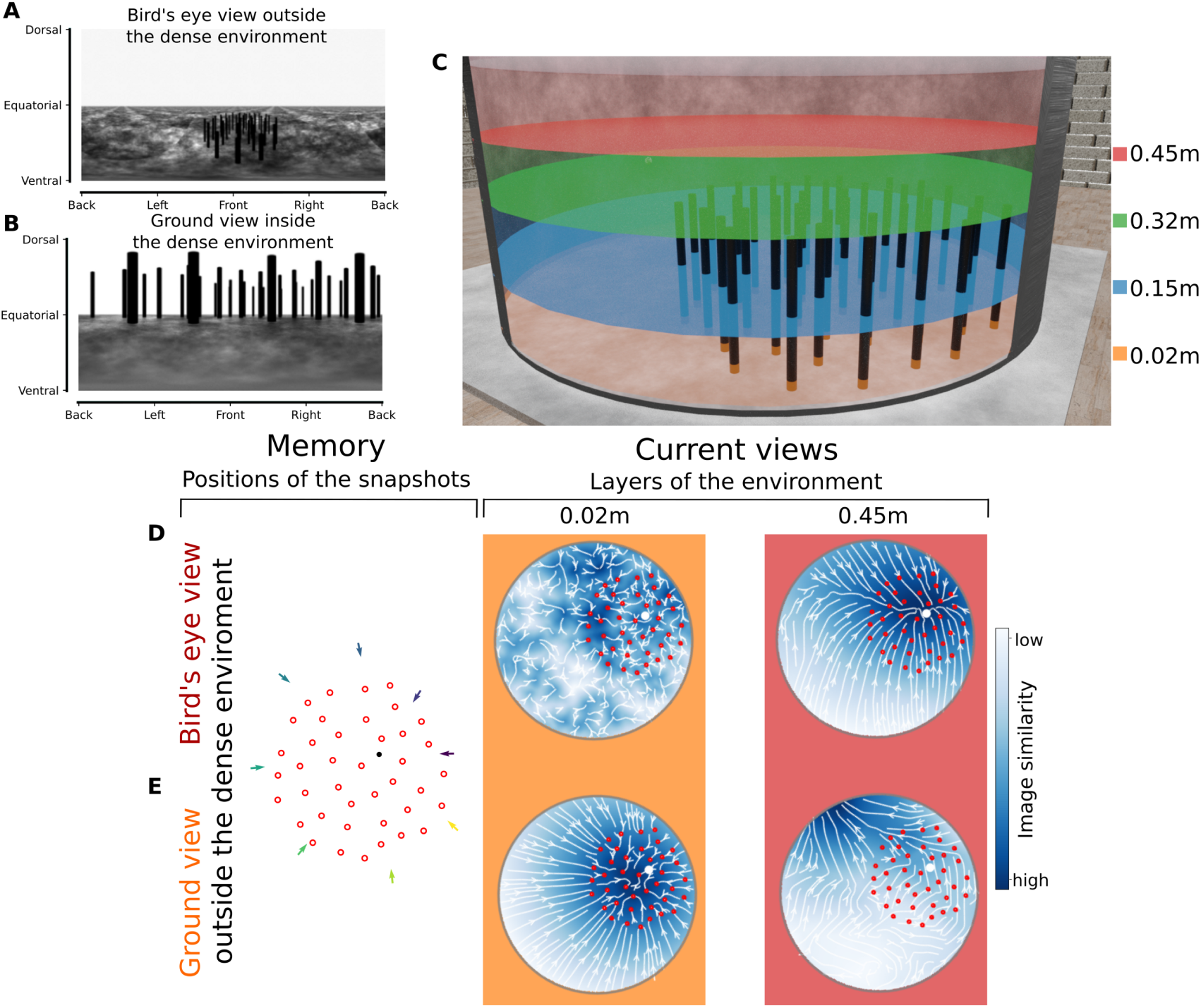
Brightness-based homing model at four altitude layers in the environment (0.02 m, 0.15 m, 0.32 m, and 0.45 m) for the area around the artificial meadow (dense environment). The rows show different parameters for the memorised snapshots (eight positions taken either outside or inside the dense environment and either above the dense environment, bird’s eye view, or close to the ground, ground views). **A&B:** Examples of panoramic snapshots in A from the bird’s eye view outside the dense environment and in B ground views inside the dense environment. The axes of the panoramic images refer to the azimuthal directions (x-axis) and to the elevational directions from the simulated bee’s point of view (y-axis, dorsal meaning upwards, ventral downwards, equatorial towards the horizon). **C:** Rendered layers of the environment for a comparison of the current view of the simulated bee. The layers are at 0.02 m(orange), 0.15 m (blue), 0.32 m(green) and 0.45 m (red) heights. **D&E:** The first column shows were the snapshots were taken in relation to the nest position (nest position in black, objects in red and snapshot positions indicated by coloured arrows). The other two columns show the comparison of memorised snapshot for two layers of the environment (0.02 m and 0.45 m as shown in C). The heatmaps show the image similarity between the current view at the position in the arena and the memorised snapshots taken around the nest (blue = very similar, white = very different). Additionally, white lines and arrows present the vector field from which the homing potential is derived. Red circles indicate the positions of the objects and the white dot indicates the nest position. The background colour of each column indicates the height of the current views that the snapshots are compared to. **D:** Memorised bird’s eye view snapshots taken outside (distance to the nest = 0.55 m) and above the dense environment (height = 0.45 m) can guide the model at the highest altitude (red background) to the nest but fails to do so at the three lower altitudes. **E:** Memorised ground view snapshots taken outside (distance to the nest = 0.55 m) the dense environment and close to the floor (height = 0.02 m) can only guide the model towards the center.

**Figure 2.**
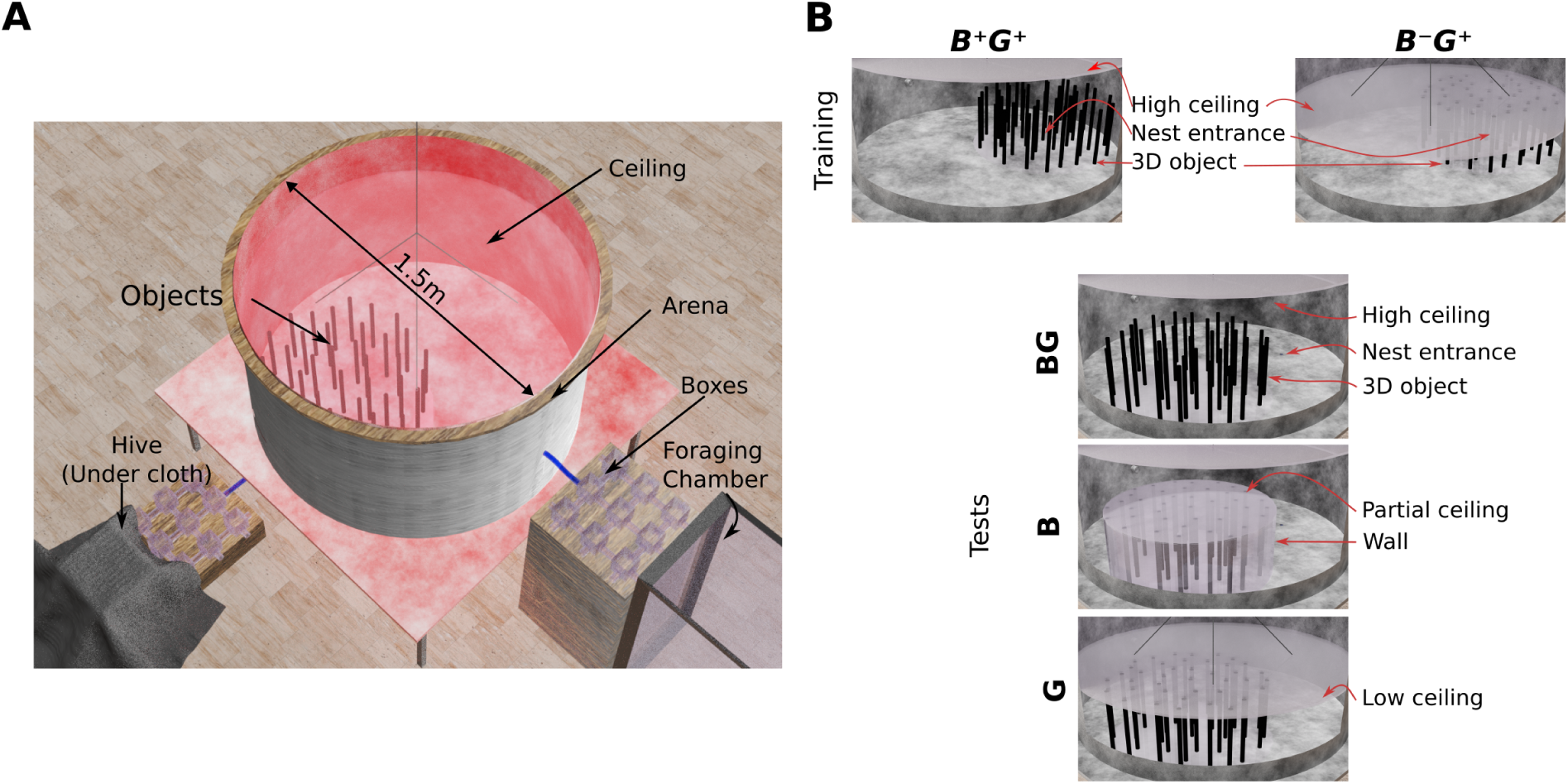
**A:** We trained two groups of bees in a cylindrical flight arena with cylindrical densely distributed objects (‘artificial meadow’) around the nest entrance. The first group, B^+^G^+^, was trained with a ceiling height of the arena twice the height of the objects providing space to fly above the objects. They might have memorised both a ground view (G^+^) as well as a bird’s eye view (B^+^) when leaving the nest. The second group, B*^−^*G^+^, was trained with an arena height restricted to the height of the objects, allowing the bees to only use ground views B*^−^*G^+^. Both groups were tested to return home in three test conditions: HighCeiling (BG), Covered (B) and LowCeilng (G). In test BG the dense environment was shifted from the training position to another position in the arena to exclude the use of potential external cues; the bees could use both, a ground view and a bird’s eye view, during return. In test B a partial ceiling above and a transparent wall was placed around the objects preventing the bees from entering the artificial dense environment during return. In test G the ceiling was lowered to the top of the objects allowing the bees to use only ground views during return. **B:** 3D view of the setup with the hive and the foraging chamber.

Eight snapshots around the nest position were compared to panoramic images either based on brightness values or contrast-weighted nearness of the rendered environment at four altitudes. These altitudes were chosen to provide information related to ecologically relevant heights for the bees during their return home (see paragraph ”Flight altitude changes with training views”). These were either close to the ground (0.02 m) resembling walking or flying close the floor, half the height of the objects (0.15 m), at the maximum height of the objects (0.32 m) or above the objects (0.45 m). We hypothesised that the simulated bee could memorise panoramic snapshots either at high altitude (bird’s eye views) or close to the floor (ground views). Further, these snapshots were taken either close to the nest (0.1 m) or just outside the dense environment (0.55 m) (Fig.1 positions of snapshots). Previous behavioural studies showed that views close to the goal are acquired during learning walks or flight [12, 44] and modelling studies could emphasise the successful use during homing [13, 15, 14]. Views just at the border of the dense environment display the change from within the dense environment to a clearance outside the dense environment. Those abrupt changes of the environment have been shown to trigger reorientation by convoluted flight patterns [9].

The comparison of simulated homing either with memorised snapshots taken from the bird’s eye or the ground views, revealed the best performance, i.e. highest image similarity and homing vectors pointing towards the nest, with bird’s eye view snapshots outside the dense environment (brightness model: Fig. 1 and Fig. S9, contrast-weighted nearness model: Fig. S13). Bird’s eye view snapshots led the simulated bee very close to the nest (brightness model: Fig. 1 D&E and Fig. S9 - S10, contrast-weighted nearness model: Fig. S13 - S14) while ground view snapshots inside the dense environment could only lead into the dense environment but not to the nest (brightness model: Fig. 1 F&G and Fig. S11 - S12, contrast-weighted nearness model: Fig. S15 and S16). Snapshots from above the dense environment showed the best homing performance when compared to images above the dense environment. Based on these results we made qualitative predictions on the bumblebees homing behaviour, assuming they are guided by homing mechanisms akin to the one tested in simulation. First, this suggests bumblebees would return to their environment by starting their approach to the goal by flying above the dense environment. Second, bees that could not acquire views above the dense environment, would not be able to locate their nest entrance.

### Ground views are sufficient for bees’ homing in a dense environment

Having shown that snapshot models perform best with bird’s eye views, we tested whether homing bees employed this strategy accordingly. Based on the model results, we hypothesised that bees should show the best homing performance when they can learn bird’s eye views and perform worse when only having access to ground views during learning. We first needed to assess their ability to return by using only visual cues provided by the dense environment. Two groups of bees were trained to find their way in a cylindrical flight arena (with a diameter of 1.5 m and a height of 0.8 m) from the nest entrance to the foraging entrance and back (Fig. 2). The group B^+^G^+^ was trained with unrestricted access to bird’s eye views (B^+^) above the dense environment and ground views (G^+^) within the dense environment. The group B*^−^*G^+^ only had access to ground views within the dense environment during the training period. To test if the bees associated the dense environment with their nest location and did not use non-intentional visual cues, e.g., outside the flight arena (even though we tried to avoid such cues), the dense environment was shifted in the tests to another position in the arena, creating two possible nest locations: the true nest location in an external reference system as during the training condition and a visual nest location relative to the dense environment (see Materials and Methods). Further, the ceiling of the arena was either placed at the height of the dense environment, allowing only ground views (G), or high above the dense environment (BG), allowing the bees to get both: ground and bird’s eye views.

In the tests BG and G where the bees had physical access to the dense environment, they searched for their nest within the dense environment. This search was in the vicinity of the visual nest entrance, though it was not always precisely centred at the nest entrance; instead, it showed some spatial spread around this location. A comparison of the time spent at the two possible nest entrance locations showed that the bees were able to find back to the visual nest in the dense environment even when the dense environment was shifted relative to an external reference frame (Fig. 3). Furthermore, we obtained a higher search percentage at the visual nest in the dense environment for the BG and G tests as well as for both groups trained to either B^+^G^+^ or B*^−^*G^+^ (Fig. 3 and Fig. S23, statistical results in SI Table 1). Most time was spent within the arena close to the visual nest location. Still, the spatial distribution of search locations also shows high search percentages in two to three other areas near the nest location (Fig. 3). These other search areas can be explained by locations similar-looking to the visual nest location between the objects. These location were also similar to those where the simulations revealed a rather broad area with a high image similarity (Fig. 1, Fig. S13 - 14 for the contrast-weighted nearness model). Both groups, B^+^G^+^ and B*^−^*G^+^, showed similar search distribution during G and BG tests, indicating that the ceiling height does not change the bees’ search distributions (t-test results with Bonferroni-correction: SI Table 1). In conclusion, when the bees could fly between the objects in the BG and G test, they were able to relocate the nest location within the dense environment by using only ground views.

**Figure 3.**
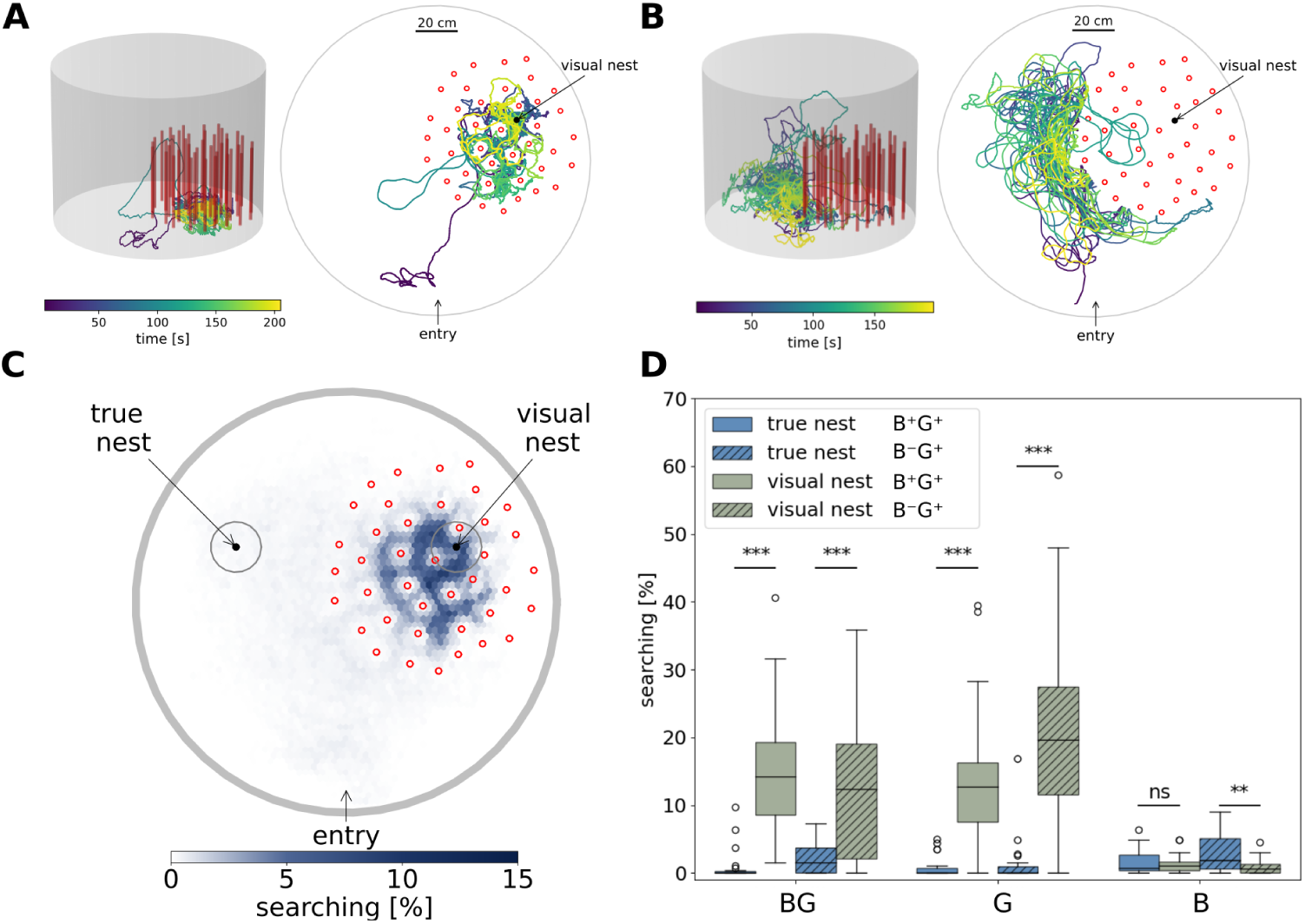
Examples of return flights and bees’ search for their nest in a dense environment (N = 26). **A&B:**Exemplary flight trajectories in 3D (left column) and a top view in 2D (right column) from the group B^+^G^+^ in the BG condition (**A**), and from the group B*^−^*G^+^ in the B condition (**B**). The colour indicates the time, blue the start and red the end of the flight. The objects are depicted by red cylinders in the 3D plot and as red circles in the 2D plot. The black dots in the 2D plot shows the visual nest position within the dense environment. **A:** The bee searches for the nest within the dense environment at a low flight altitude. **B:** The bee is mainly trying to enter the covered dense environment from the side. **C:** Spatial search distribution represented by hexagonal binning of the percentage of visits of all bees (relative to each bee’s total flight time) in the BG condition from group B^+^G^+^. Orange circles indicate the true and the visual nest position. Black circles indicate the object positions of the dense environment. **D:** Percentage searching for the two groups B^+^G^+^ (filled boxes, N = 26) and B*^−^*G^+^ (hatched boxes, N = 26) in the tests BF, G and B relative to the total flight time. The search percentage at the true nest is given in blue and at the visual nest in green. For all tested conditions and both groups, the bees searched more at the visual nest within the dense environment than at the true nest location (refer to SI Table 1 for statistical tests).

### Bird’s eye views are not sufficient for bees to return

Based on the results of the simulations (Fig. 1), we hypothesised that bees could pinpoint their nest position by using only bird’s eye views. In addition, we observed that during the first outbound flights, bees quickly increased their altitude to fly above objects surrounding the nest entrance (Fig. S22). Therefore, we restricted the bees’ access to the dense environment during their return flight using a transparent cover around it, so they did not have access to the dense environment from the side but only from above (see test B, Fig. 2 A). In this test, the bees tried to enter the dense environment sideways between the objects as the search distributions show (Fig. 5 B&D). The bees did not search at the nest location above the dense environment (Fig. 5 B&D) which they could have done if they would have learned to enter the dense environment from above. We can therefore reject the hypothesis that bees use bird’s eye view to return home in a dense environment. Rather ground views seem to be sufficient for homing. Nevertheless, an interesting aspect with respect to finding the nest hole is revealed by the entry positions into the dense environment. Entry points from all three tests (B, BG and G for both groups) from the top and the side into the dense environment supported the finding that most bees tried to cross the boundary to the dense environment from the side while only very few tried to enter the dense environment from the top (Fig. 4). The entry points from the side concentrate mainly on three locations indicating that the bees learned similar entrance points to the dense environment (Fig. 4 A). Taken together, these results show that the bees learned certain entry points of the dense environment. This learning could have taken place during learning flight, if the bees exited the dense environment at these locations predominantly, or learning taking place following return trials, or both.

**Figure 4.**
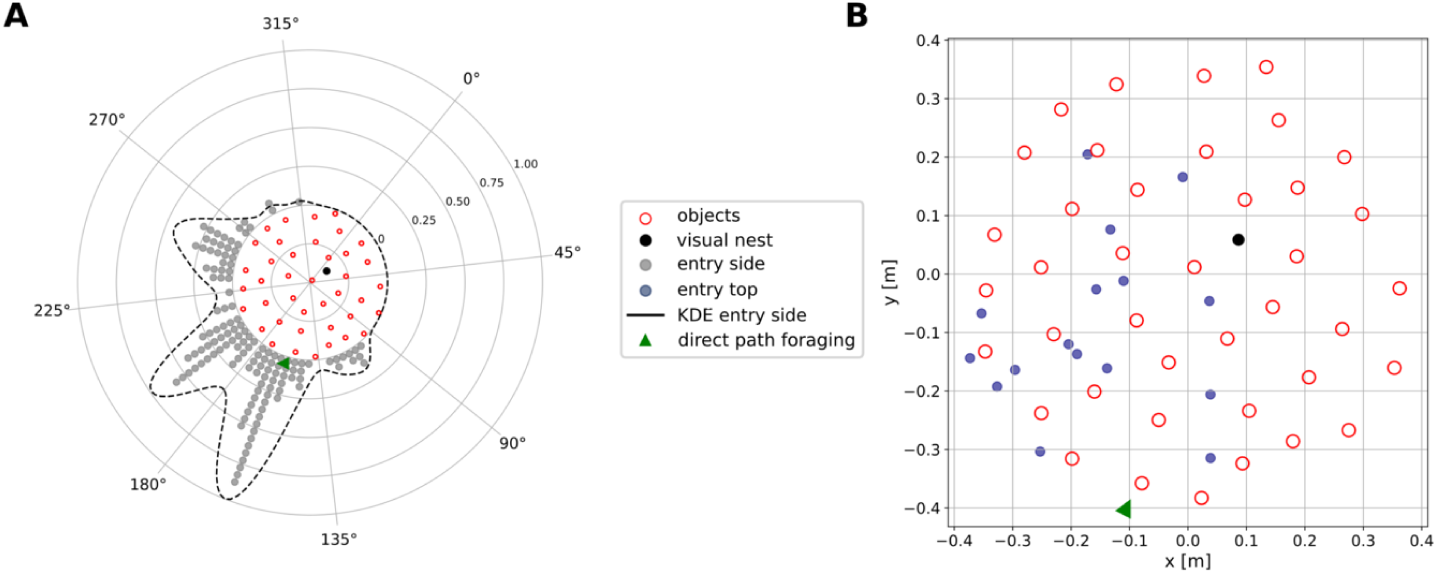
Entry points of the bees of both groups (N = 52) in all three tests (total entry points = 149) to the dense environment, from the side (**A**, circular histogram in grey) and from the top (**B**, scatter plot in blue) of the dense environment. The direction of the direct path from the arena entrance to the nest is given by the green triangle. The kernel density estimation (KDE) of the entries from the side of the dense environment is shown as a black, dashed line in **A**. The radial axes represents the normalized magnitude of the KDE.

**Figure 5.**
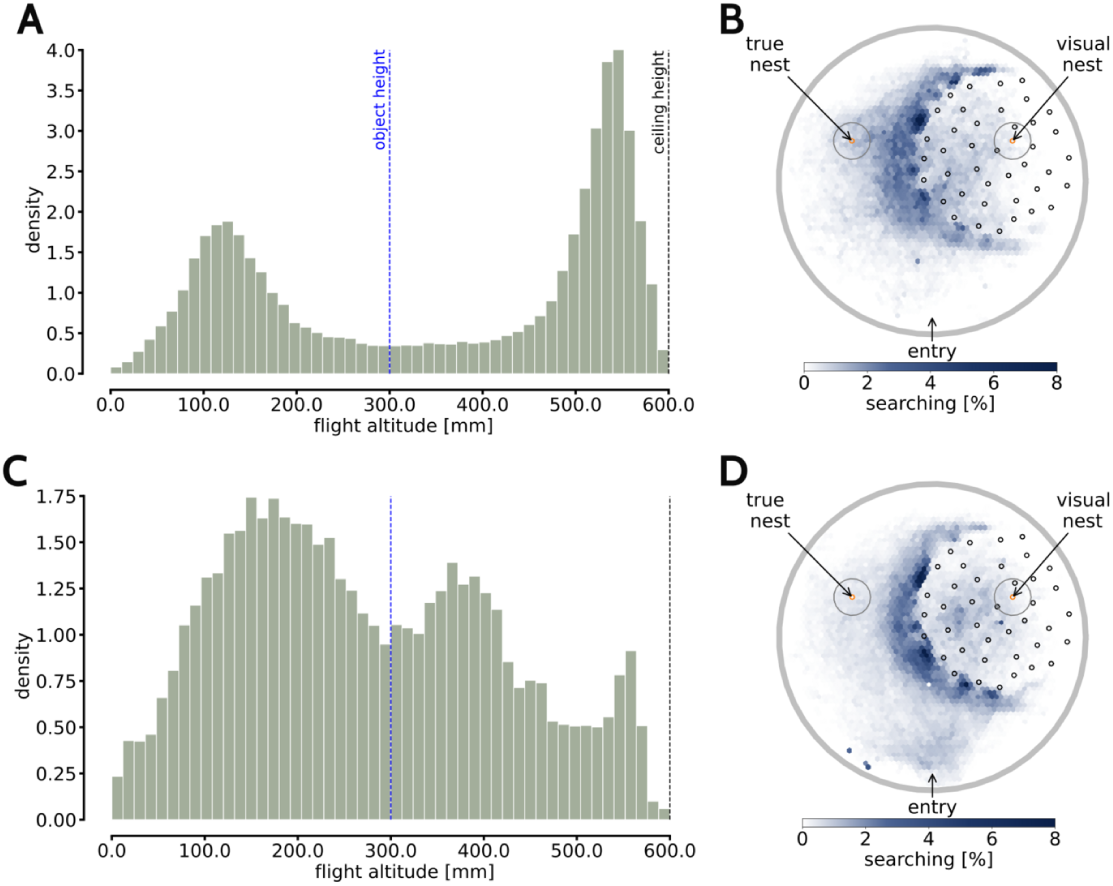
Probability density distribution of the flight altitude and the search distribution for the B condition of the groups B*^−^*G^+^ (**A&B**) and B^+^G^+^ **C&D**). A flight altitude of 0 is at the floor, of 300mm is at the height of the objects and 600mm at the ceiling. **A**: The group B*^−^*G^+^ constrained to a low ceiling during training shows two peaks for a low altitude and for a high altitude. **C**: The group B^+^G^+^ shows a broader distribution, with three peaks either half the height of the objects, just above the height of the objects, or close below the ceiling. **B&D**: The search distributions reveal that the bees tried to enter the dense environment between the objects.

### Flight altitude changes with training views

We investigated the influence of the potentially available views during training on the bees’ altitude during homing by comparing the employed flight altitudes. The flight altitude probability density distributions for the G and BG tests showed for both groups that the bees flew mostly very low searching for the nest entrance (Fig. S24). The distributions, however, looked very different for the test B. The flight altitude distribution of the group B*^−^*G^+^ shows two peaks, one around 130mm and one around 550mm (Fig. 5 A). These peaks indicate that the bees either flew just below half the object height, probably trying to enter the covered dense environment from the side or they flew close to the ceiling of the arena. The flights close to the ceiling might reflect exploratory behaviour in a section of the flight arena the bees could not experience before as they were trained to a ceiling constrained to the height of the objects. The behaviour of flying at a low altitude, trying to enter the dense environment from the side, is fitting to the peaks of search time around the covered dense environment (Fig. 2 A&B). The group B^+^G^+^ showed a broader probability density distribution indicating three shallower peaks, one at an altitude around 170mm, which refers to half the height of the objects, a second peak around 370mm, referring to just above the objects and a third peak just below the ceiling where the bees might have explored the boundaries of the arena. This indicates that the bees trained with B^+^G^+^ seemed to have learned to fly above the objects and used this information when the cover blocked the entrance to the dense environment. The group B*^−^*G^+^, constrained to fly only up to the top of the objects but not above, seemed to be unable to search for another way to enter the dense environment from above. Although the bees exposed to B^+^G^+^ during training, did not search above the nest position, they seemed to have learned the height of the objects which the altitude-constrained group, B*^−^*G^+^, could not learn because the bees flew more at the height of the objects. Overall, there is a clear difference in the flight altitude distributions between the two groups and between the tests. The group trained with B^+^G^+^ flew more at the height of the objects when the dense environment was covered, while the group trained with a B*^−^*G^+^ seemed to just explore the newly available space at a very high altitude with flying little at the height of the objects. This indicates that the flight altitude during learning plays a role in what altitude the bees fly during the return flights, and that this could influence what aspects are learned.

## Discussion

We investigated how bumblebees find home in a dense environment using views at different altitudes. The results of simulations of nest-centered snapshot-based homing models [53] suggested the best homing performance with bird’s eye views from above the dense constellation of objects. Surprisingly, bees performed equally well whether or not they could experience bird’s eye views during training or just ground views, i.e. views taken close to the ground. Even bees trained with both ground and bird’s eye views predominantly did not use bird’s eye views, because they usually flew into the dense environment from the side and not from above. Thus, instead of solely relying on snapshots, we hypothesise a multi-step process for homing in a dense environment relying on a variety of tools from the navigational toolkit supported by the bees’ behaviour.

### Working range of snapshot models is limited inside a dense environment

Snapshot matching is widely assumed to be one of the most prominent navigational strategies of central-place foraging insects [52]. In the simulations of this study, we found that bird’s eye view snapshots led to the best homing performance, while ground view snapshots could only find the dense environment but not the nest position within the dense environment. Additionally, the model steered only to the nest when the current views were from above the objects (altitudes 0.32 m and 0.45 m). A previous study showed that snapshots taken at higher altitudes result in larger catchment areas than snapshots taken close to the ground [33]. Aligned with this study, we found that image difference functions pointed towards the location of the objects surrounding the nest when the images were taken above the objects. However, within the dense set of objects surrounding the nest, the model did not perform well in pinpointing the nest position (Fig. 1D&E). Furthermore, in a particular form of snapshot model, i.e. skyline snapshots, occluding objects lead to smaller catchment areas [34]. Our model simulations support both conclusions, as the bird’s eye views from above the dense environment were less occluded. Behavioural studies also confirm the advantage of higher altitude snapshots as it was found that honeybees and bumblebees may use ground features of the environment while navigating on a large spatial scale (i.e. of hundreds of metres) [31, 3]. A modelling and robotic approach of Stankiewicz and Webb [42] could also confirm these findings. Overall, our model analysis could show that snapshot models are not able to find home with views within a dense environment but only with views from above it.

### Bees’ homing performance in a dense environment

We found that the behavioural performance of bumblebees differed quite substantially from our snapshot-based model predictions, when we tested bees either trained with bird’s and ground views or trained only with ground views. Both groups of bees performed equally well in finding the nest position between the densely distributed objects. This finding suggests that even when bees had the opportunity to fly above the objects of the dense environment during training, they relied on means other than bird’s eye views to find their goal. From a flight control perspective, changing between flying up and down could be energetically more demanding than between left and right. Large-scale studies (e.g. [31]) state that bees fly more in a plane than up/down. Moreover, from a flight control perspective, it might be difficult for the bees to approach the dense environment from above and then descend to the nest between the objects. This hypothesis is supported by a study showing that bees prefer to avoid flying above obstacles at short distances when they have the choice to fly around [46].

### Hypothetical mechanisms of homing in dense environments

We observed during the first outbound flights of four recorded bees that bees flew above the clutter (for a systematic analysis of the outbound flights see [41]). This behaviour suggests that bees may acquire nest-centered snapshots during these flights, allowing them to return home, as supported by our model simulations (Fig. 1) based on an extensively tested model [54]. However, we observed that bees typically return to their nest by flying below object height and navigating within the dense environment. Since we did not record the bees’ full visual history—being beyond the study’s scope and only becoming relevant due to inconsistencies between modeling and behavioral results—we can only hypothesize a possible mechanism by which experienced bees return to their nest based on known navigation mechanisms. These hypotheses are grounded in known navigation mechanisms such as snapshot-based route following and usage of a home vector.

In snapshot-based route following models, the animal compares its current view with stored snapshots of a route[2]. For successful navigation, the animal must collect sufficient visual memories along the route. Indeed, as the animal distances itself from the memorized view, it may be led astray. In other terms, each memorised snapshot allows the animal to follow the route within a given area. The set of memorised snapshots forms a catchment area, defined similar to a funnel around the route [2]. The challenge to build such a memory is at least twofold. First, the catchment area of a single snapshot decreases with increasing clutter [34]. This would require a bee to memorised many snapshots (the neural capacity of bees has been estimated with model simulation to 100,000 snapshots [1]). Second, the snapshots need to be either acquired along the route [2] or associated to attractive (resp. repulsive) weights [26]. Due to the high number of objects in clutter, it might be challenging to build a set of route-aligned snapshots.

The B^+^G^+^ bees (those exposed to both ground and bird’s eye views) might overcome these challenges by utilizing a two-step strategy. During their initial outbound flights, they could use snapshots taken from above the clutter, which require fewer images to be effective. Then, through a trial-and-error approach, the bees could attempt to return from within the dense environment, resorting to the higher-altitude snapshots if they failed. This strategy would be effective for B^+^G^+^ bees because during training they had access to both the cluttered and open regions. In contrast, the B*^−^*G^+^ bees, which were only exposed to ground views, also managed to find their way back, indicating they might have relied on a different navigational mechanism.

Since the route-following model does not fully explain the behaviour of the bees, we propose an alternative: visual memories could trigger a reloading of a home vector, as described by Webb [47]. This hypothesis suggests that bees can reload this vector when stimulated by visual cues. In our study, the memorised home vector, obtained by mechanisms like path integration or others, would guide a bee to the nest based on its initial position (Fig. 6, black arrow). In the training setup, the bee’s home vector and visual cues, such as the densely distributed objects around the nest, were congruent (Fig. 6, green arrow). In the test setup, the visual nest hole was displaced, but the bees likely relied on the dense environment as a salient visual cue for homing when entering the flight arena from the foraging chamber. Bees may use this dense object constellation to associate it with their nest, which could then be found by a relatively straight path (Fig. 6, green arrow), supported by the high concentration of entry points near the dense environment (Fig. 4). These preferred entry positions used by bees suggest that they memorised specific locations to enter the dense environment, possibly triggered by visual cues like snapshots at the boundary (Fig. 6, yellow arrow). These visual memories might help refine the bee’s vector memory and guide them along the environment’s border until they find a familiar entry point. Once inside, bees could use the distance between the nest and the wall, along with their memorised direction, to locate the nest 6, blue arrow). This multi-faceted navigation strategy, which integrates visual memories, environmental landmarks, and vector-based navigation, underscores the complexity and flexibility of bees’ homing behaviour in dense environments and may serve as a working hypothesis for future studies.

**Figure 6.**
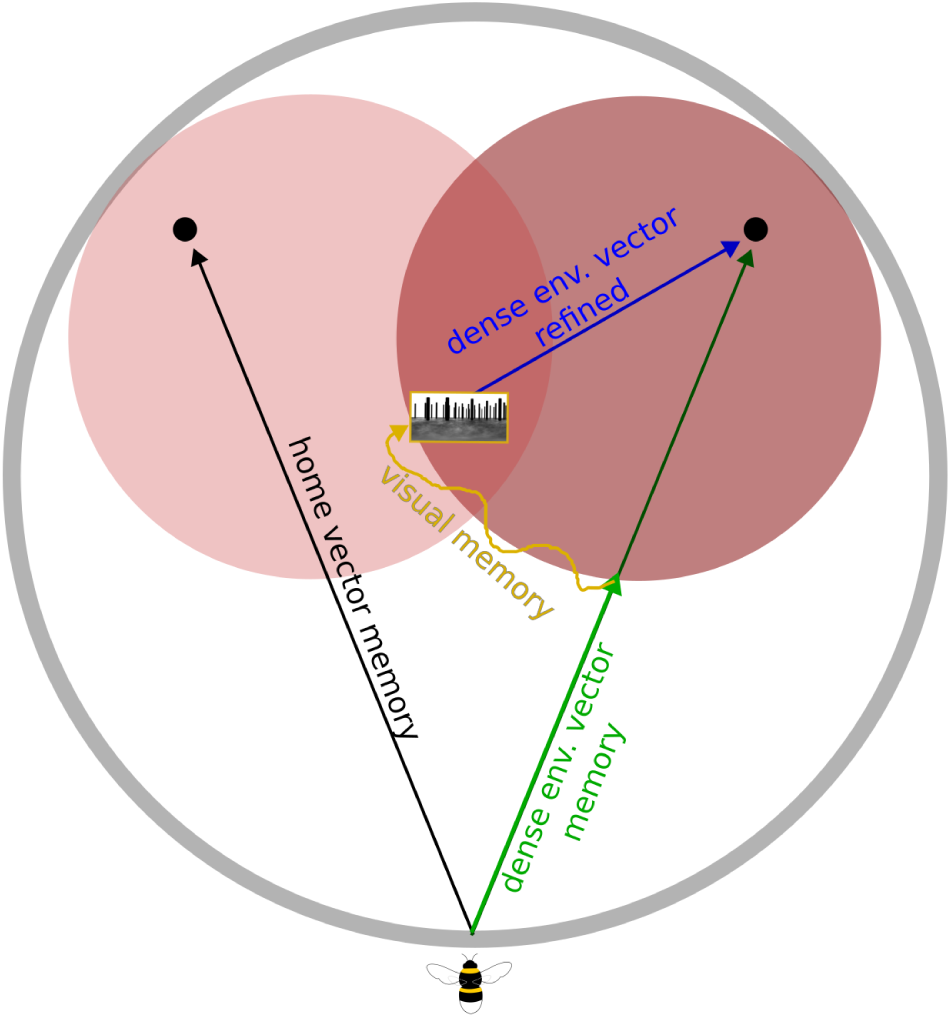
Schematic of a hypothetical homing mechanisms based on multiple navigational strategies. A bee entering the arena (grey circle) could have memorised a home vector (black arrow) pointing towards the true nest location at the training position in the dense environment (black dot within the light red circle). However, the visual scene changed drastically in the test. Hence, a vector memory might point coarsely to the dense environment (light green arrow) triggered by the prominent visual cue of the dense environment shifted to the test position (dark red circle). While some bees might use this vector providing the shortest path to the nest, other bees could rely on visual memories for entering the clutter. They will search at the border of the dense environment (curved, yellow arrow) for a previously experienced entry position to the dense environment (visual memory, framed in yellow). This position could reload a refined dense environment vector pointing to the more precise nest position within the dense environment (blue arrow).

### Outlook

Our study on 3D homing of bees underscores the limitations inherent in current snapshot models, revealing their inability to provide precise positional estimates within dense environments, especially when compared to the navigational abilities of bees. Notably, bees that had the opportunity to learn during training bird’s-eye views demonstrated adaptability in spatially constrained situations, although this strategy was not employed explicitly for searching nests above the dense environment. Having shown that bumblebees are capable of finding their nest hole in very challenging, dense environments, we investigated the influence of the environmental structure of the nest surroundings on the learning flights of bumblebees in a separate study [41]. Future research is needed to extend our findings on a larger scale and explore the development of 3D snapshot models that account for altitude variations. Furthermore, these insights extend beyond bees and may have implications for other species, like birds with altitude fluctuations during foraging or nesting [4]. The use of bird’s-eye views in dense forests, akin to our findings with bees, prompts consideration of similar behaviours in other flying animals, but also for walking animals, such as ants such navigating varied vegetation heights [18]. Switching views might considerably affect an animal’s ability to solve spatial problems, as shown in humans [22]. Understanding how best to combine this information for navigation will benefit the development of autonomously flying robots—for instance, helping them navigate in dense environments.

## Materials and Methods

### View-based homing models

We rendered an environment with 40 randomly placed red cylinders, creating an artificial meadow (dense environment), surrounding the nest entrance placed on a floor with a black and white pattern with a 1/f spatial frequency distribution (a distribution observed in nature [40]) in a graphic software (Blender, 2.82a). A panoramic image was taken along a grid with 0.5 cm steps within the dense environment and 1 cm steps in the remaing part of the arena. The arena was 1.5 m in diameter like for the following behavioural experiments. For the rendering we focused only on relevant cues like the objects and the patterned floor. We did not render the nest hole which was visually covered during the behavioural experiments. In addition, the arena wall was also not rendered as it did not contain any information for bees other than for flight control. Further simulations with a rendered arena wall led to worse results because the agent was mainly led to the center of the arena (Fig. S17, Fig. S18-21). Therefore, we focused our analysis on the simulation with a rendered arena wall. The grid-spacing was used at four altitudes: 2 cm for walking or hovering bees, 15 cm for bees flying half of the object height, 32 cm for bees flying just above the objects and 45 cm for bees flying very high to compare how the calculated similarity changes regarding the altitude. We expected qualitatively similar results for analyses with different volumes or catchment areas. In the next step, eight images around the nest were taken and stored as memorised snapshots, and the current views were compared (examples in Fig. 7-8).

**Figure 7.**
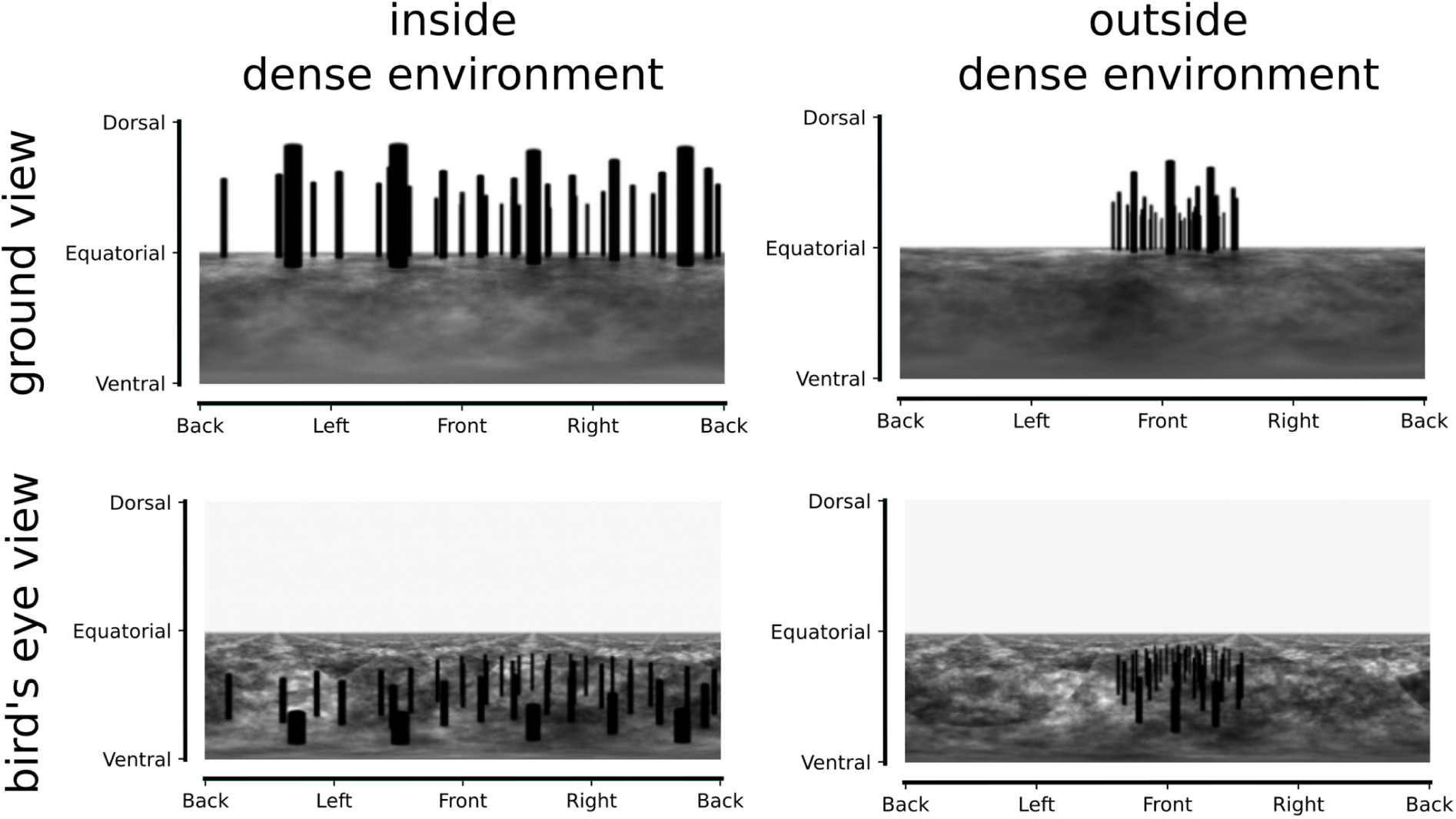
Examples of the memorised snapshots based on brightness values for inside and outside the dense environment as well as from the bird’s and ground view perspective. The axes of the panoramic images refer to the azimuthal directions (x-axis) and to the elevational directions from the simulated bee’s point of view (y-axis, dorsal meaning upwards, ventral downwards, equatorial towards the horizon).

**Figure 8.**
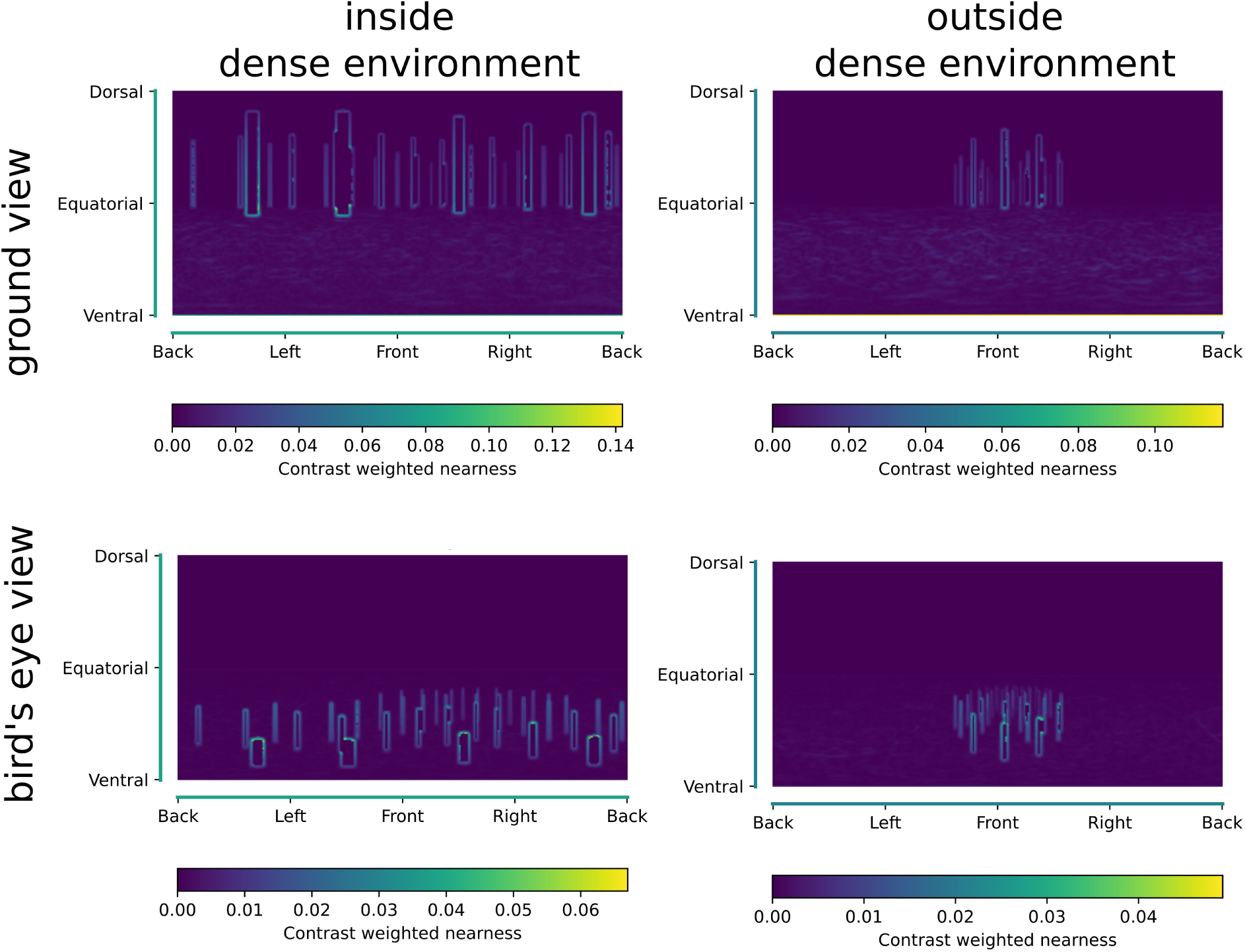
Examples of the memorised snapshots based on contrast-weighted nearness for inside and outside the dense environment as well as from the bird’s and ground view perspective. The axes of the panoramic images refer to the azimuthal directions (x-axis) and to the elevational directions from the simulated bee’s point of view (y-axis, dorsal meaning upwards, ventral downwards, equatorial towards the horizon).

**Figure 9.**
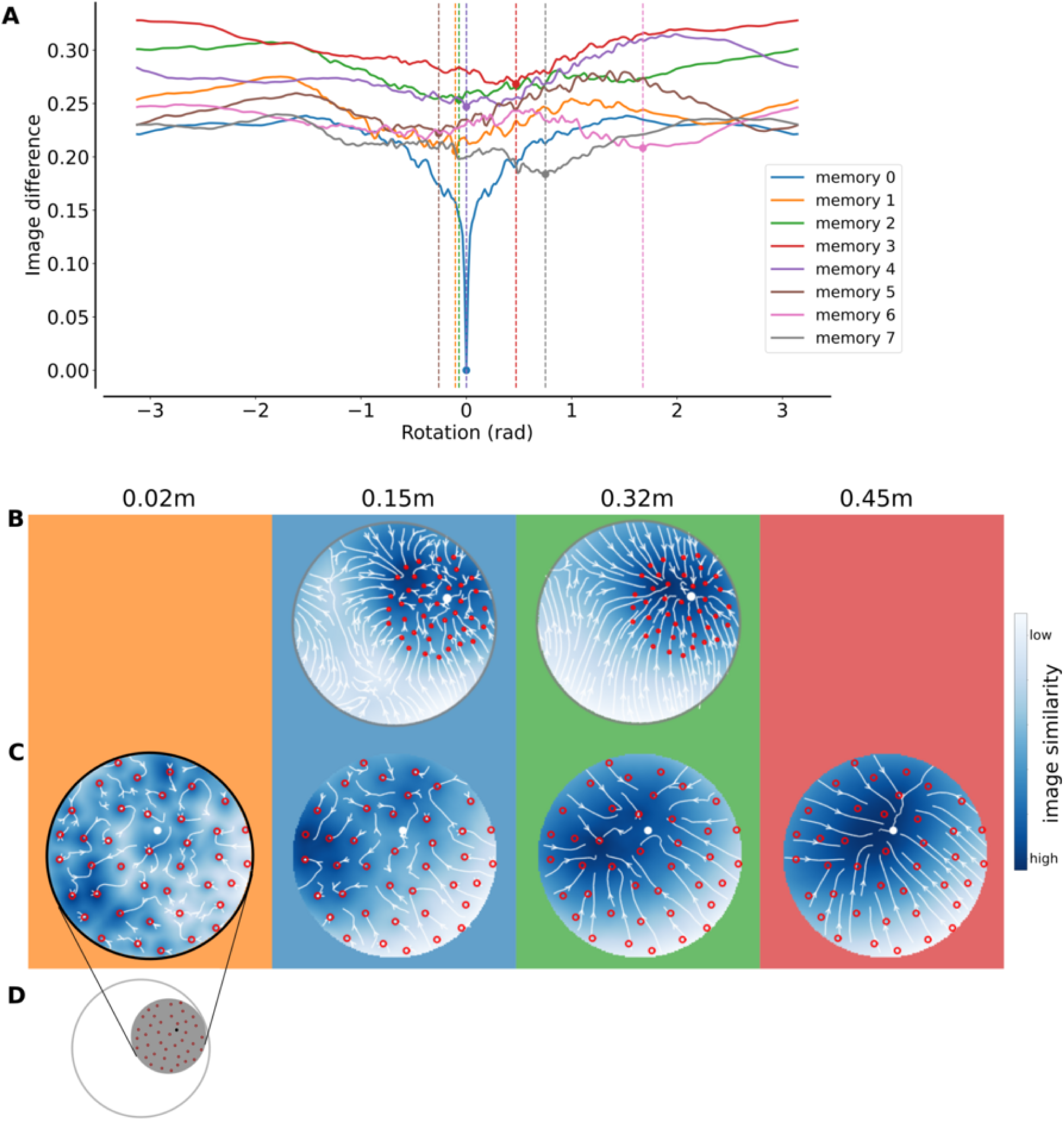
Brightness-based homing model at four altitudes (0.02 m, 0.15 m, 0.32 m, and 0.45 m) with bird’s eye view snapshots taken outside the dense environment (distance = 0.55 m, height = 0.45 m). **A:** We applied the model to a list of images, these images are examples of the eight memorised snapshots which are aligned in the nest direction and taken around the nest. For the rotational image difference function, we used memory 0 as a reference image, and compared the seven others by rotating them against memory 0. We observe that the image difference function is minimum for the memorised image at a null rotation, as expected. If the other images are not two far from the nest, we may see other local minima for each of these images, where the local minima are shifted according the nest bearing. **B-C:** Heatmaps of full environment (**B**) and only the dense area (**C**, as shown in **D**) of the image similarity. The colour indicates the image difference between the view at the position in the arena and the snapshots taken around the nest. The model leads the simulated bees with bird’s eye view snapshots outside the dense environment to the nest position when the images are compared to images at an altitude of 0.45 m.

**Figure 10.**
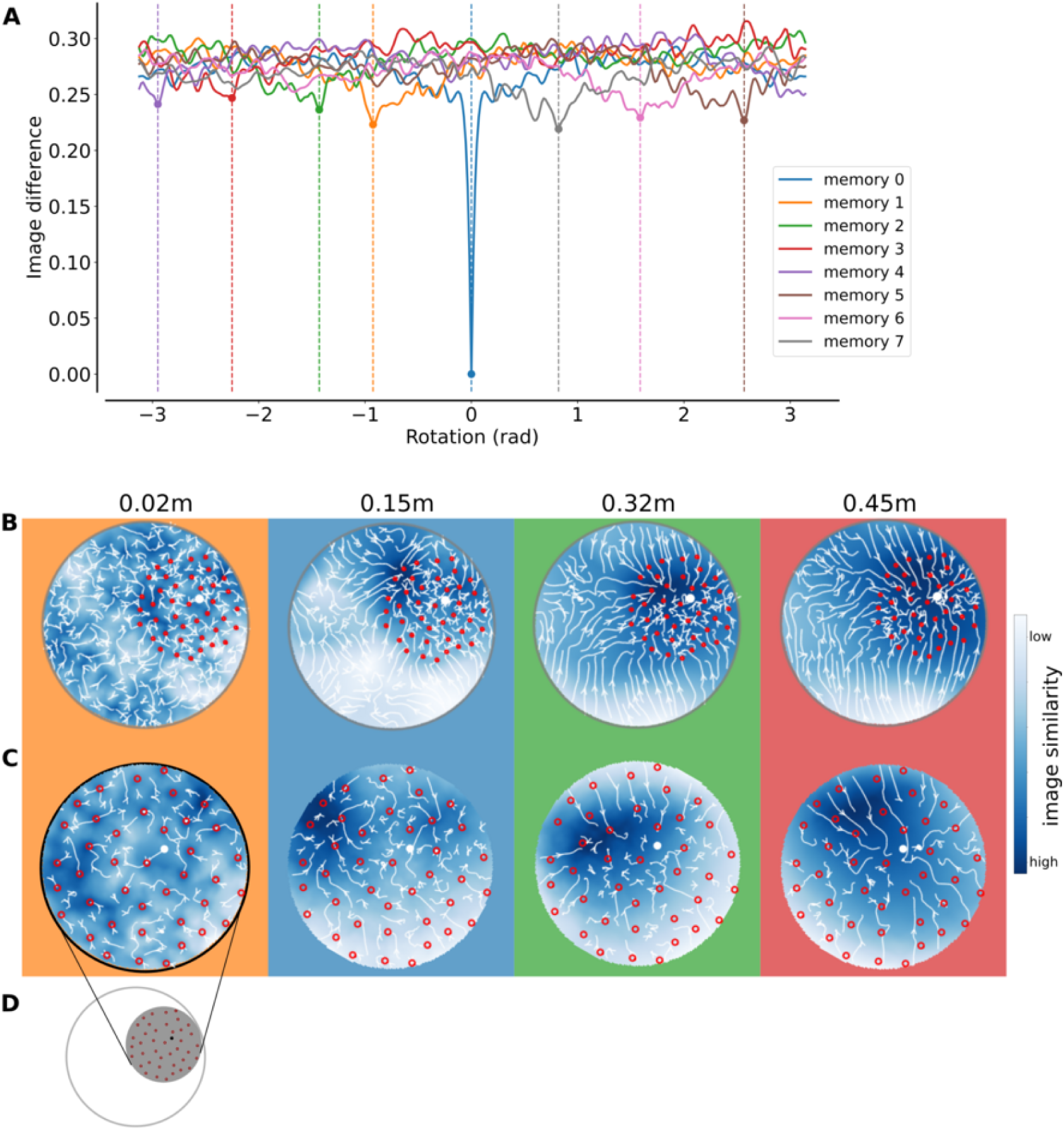
Brightness-based homing model at four altitudes (0.02 m, 0.15 m, 0.32 m, and 0.45 m) with bird’s eye view snapshots taken inside the dense environment (distance = 0.1 m, height = 0.45 m). The colour indicates the image difference between the view at the position in the arena and the snapshots taken around the nest. **A:** We applied the model to a list of images, these images are examples of the eight memorised snapshots which are aligned in the nest direction and taken around the nest. For the rotational image difference function, we used memory 0 as a reference image, and compared the seven others by rotating them against memory 0. We observe that the image difference function is minimum for the memorised image at a null rotation, as expected. If the other images are not two far from the nest, we may see other local minima for each of these images, where the local minima are shifted according the nest bearing. **B:** Heatmaps of full environment (**B**) and only the dense area (**C**, as shown in **D**) of the image similarity. The colour indicates the image difference between the view at the position in the arena and the snapshots taken around the nest. The model leads the simulated bees, with bird’s eye view snapshots inside the dense environment, only coarsely to the dense environment but not to the nest when the images are compared to images at an altitude of 0.45 m.

**Figure 11.**
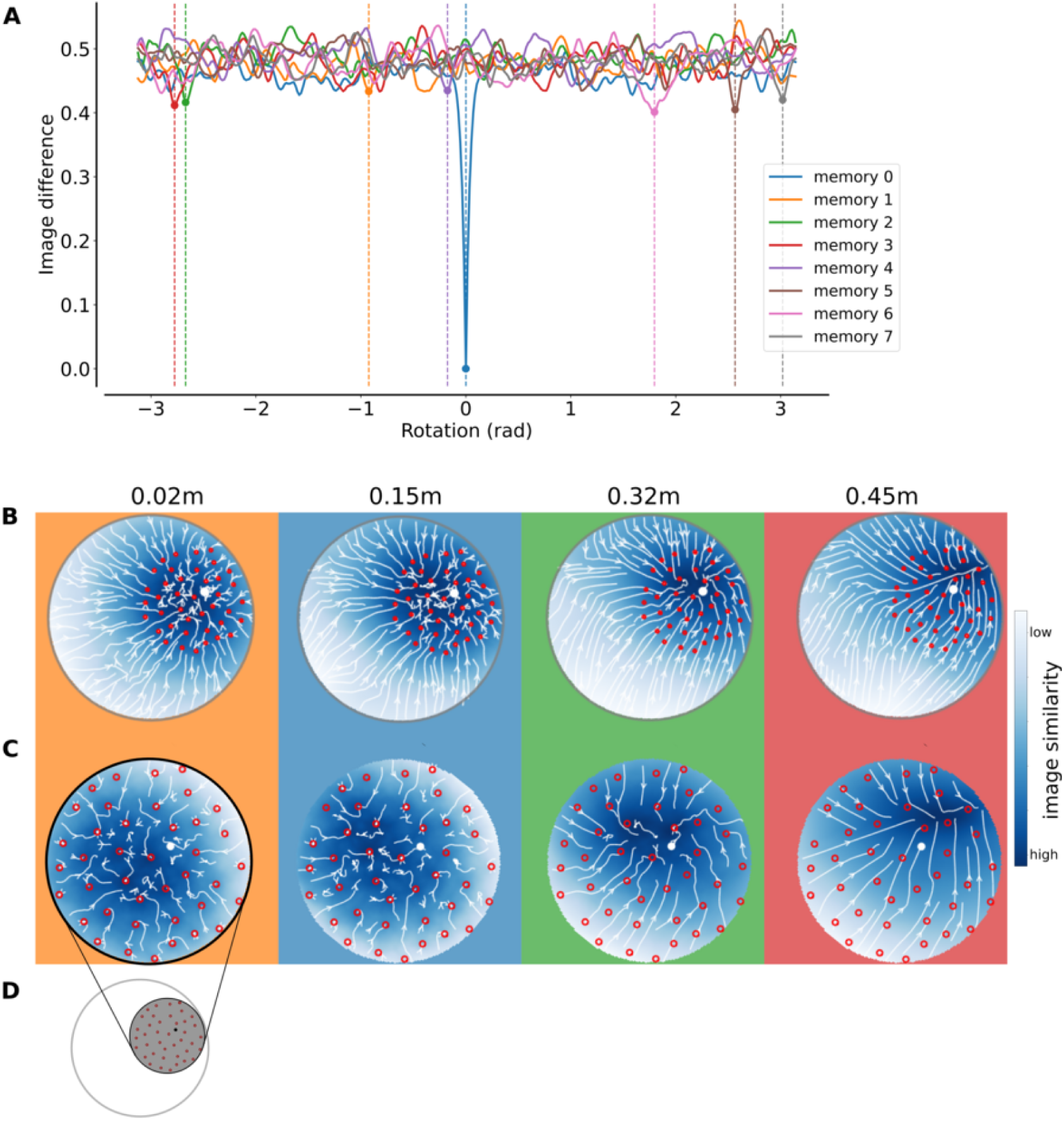
Brightness-based homing model at four altitudes (0.02 m, 0.15 m, 0.32 m, and 0.45 m) with ground view snapshots taken inside the dense environment (distance = 0.1 m, height = 0.02 m). The colour indicates the image difference between the view at the position in the arena and the snapshots taken around the nest. **A:** We applied the model to a list of images, these images are examples of the eight memorised snapshots which are aligned in the nest direction and taken around the nest. For the rotational image difference function, we used memory 0 as a reference image, and compared the seven others by rotating them against memory 0. We observe that the image difference function is minimum for the memorised image at a null rotation, as expected. If the other images are not two far from the nest, we may see other local minima for each of these images, where the local minima are shifted according the nest bearing. **B:** Heatmaps of full environment (**B**) and only the dense area (**C**, as shown in **D**) of the image similarity. The colour indicates the image difference between the view at the position in the arena and the snapshots taken around the nest. The model leads the simulated bees, with ground view snapshots inside the dense environment, only to the center of the dense environment (altitude of 0.32 m) or shifted away from the nest (altitude of 0.45 m) but not to the nest.

**Figure 12.**
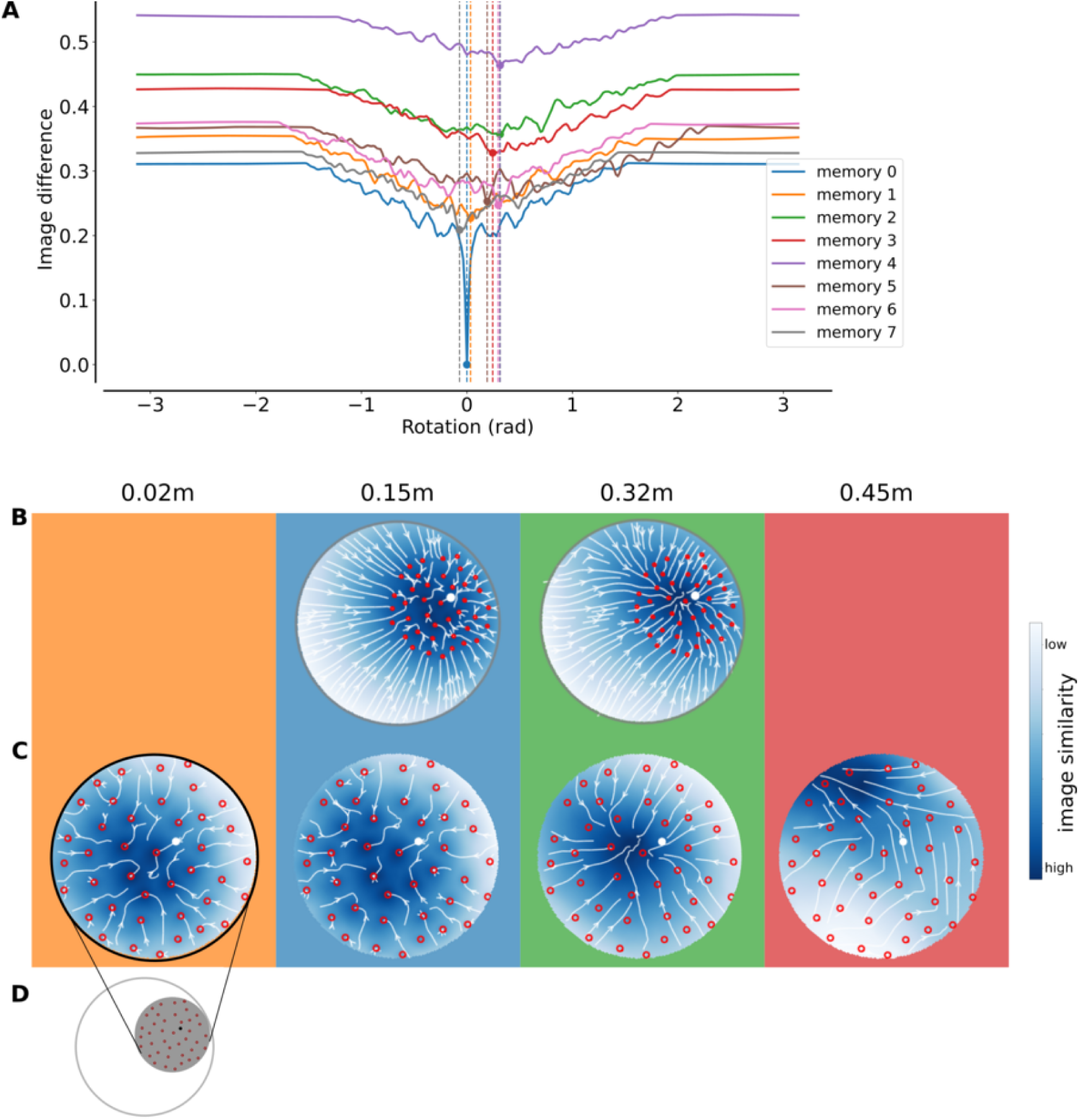
Brightness-based homing model at four altitudes (0.02 m, 0.15 m, 0.32 m, and 0.45 m) with ground view snapshots taken outside the dense environment (distance = 0.55 m, height = 0.02 m). The colour indicates the image difference between the view at the position in the arena and the snapshots taken around the nest. **A:** We applied the model to a list of images, these images are examples of the eight memorised snapshots which are aligned in the nest direction and taken around the nest. For the rotational image difference function, we used memory 0 as a reference image, and compared the seven others by rotating them against memory 0. We observe that the image difference function is minimum for the memorised image at a null rotation, as expected. If the other images are not two far from the nest, we may see other local minima for each of these image, where the local minima are shifted according the nest bearing. **B:** Heatmaps of full environment (**B**) and only the dense area (**C**, as shown in **D**) of the image similarity. The colour indicates the image difference between the view at the position in the arena and the snapshots taken around the nest. The model leads the simulated bees, with ground view snapshots outside the dense environment, only to the center of the dense environment (altitude of 0.32 m) but not to the nest.

**Figure 13.**
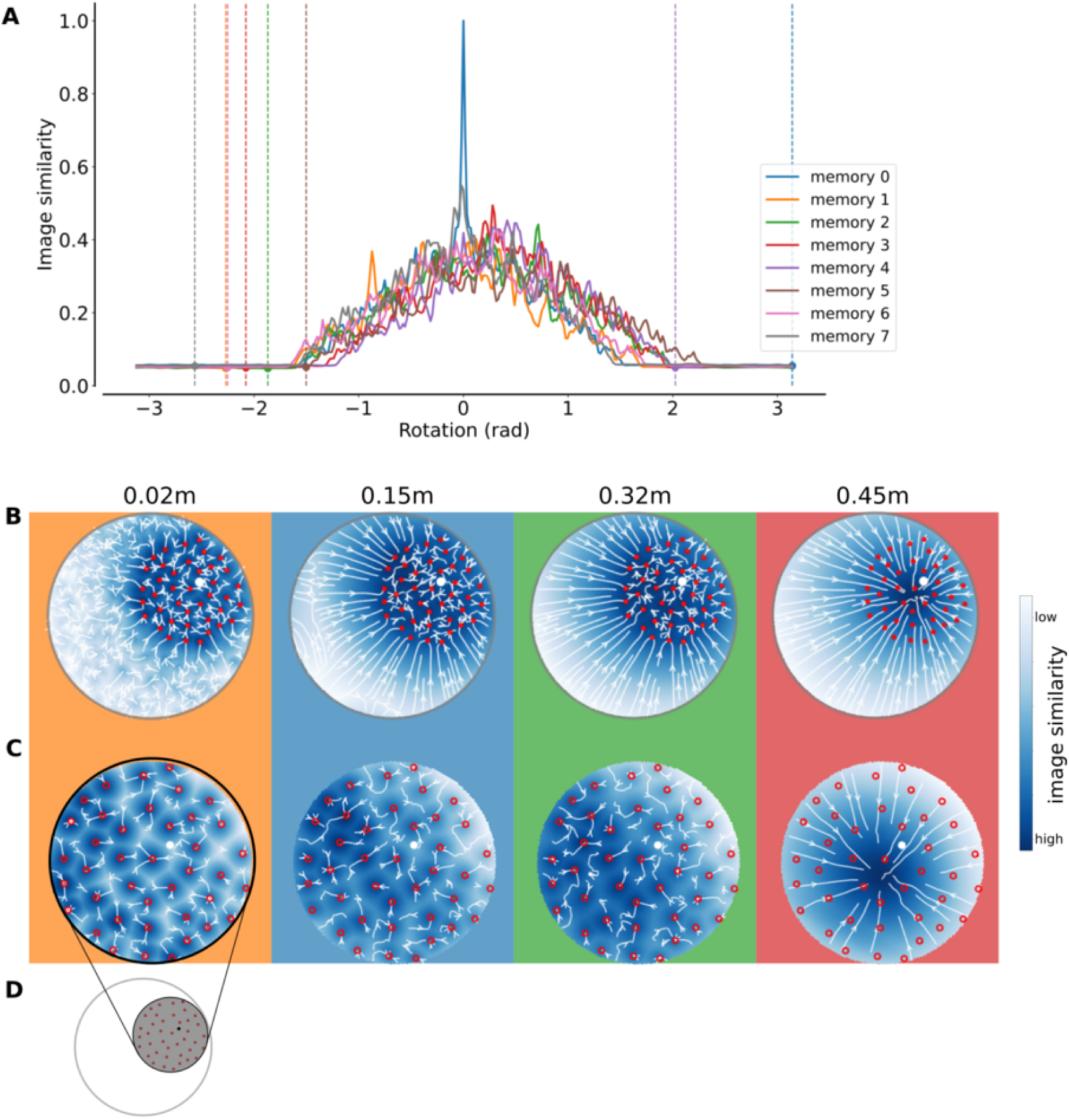
Contrast-weighted nearness based homing model at four altitudes (0.02 m, 0.15 m, 0.32 m, and 0.45 m) with bird’s eye snapshots taken outside the dense environment (distance = 0.55 m, height = 0.45 m). **A:** We applied the model to a list of images, these images are examples of the eight memorised snapshots which are aligned in the nest direction and taken around the nest. For the rotational image difference function, we used memory 0 as a reference image, and compared the seven others by rotating them against memory 0. We observe that the image similarity as based on [14] is the maximum for the memorised image at a null rotation, as expected. If the other images are not two far from the nest, we may see other local maxima for each of these image, where the local maxima are shifted according the nest bearing. **B:** Heatmaps of full environment (**B**) and only the dense area (**C**, as shown in **D**) of the image similarity. The colour indicates the image similarity between the view at the position in the arena and the snapshots taken around the nest. The model leads the simulated bees, with bird’s eye view snapshots outside the dense environment, only to the center of the dense environment (altitude of 0.32 m) or shifted away from the nest (altitude of 0.45 m) but not to the nest.

**Figure 14.**
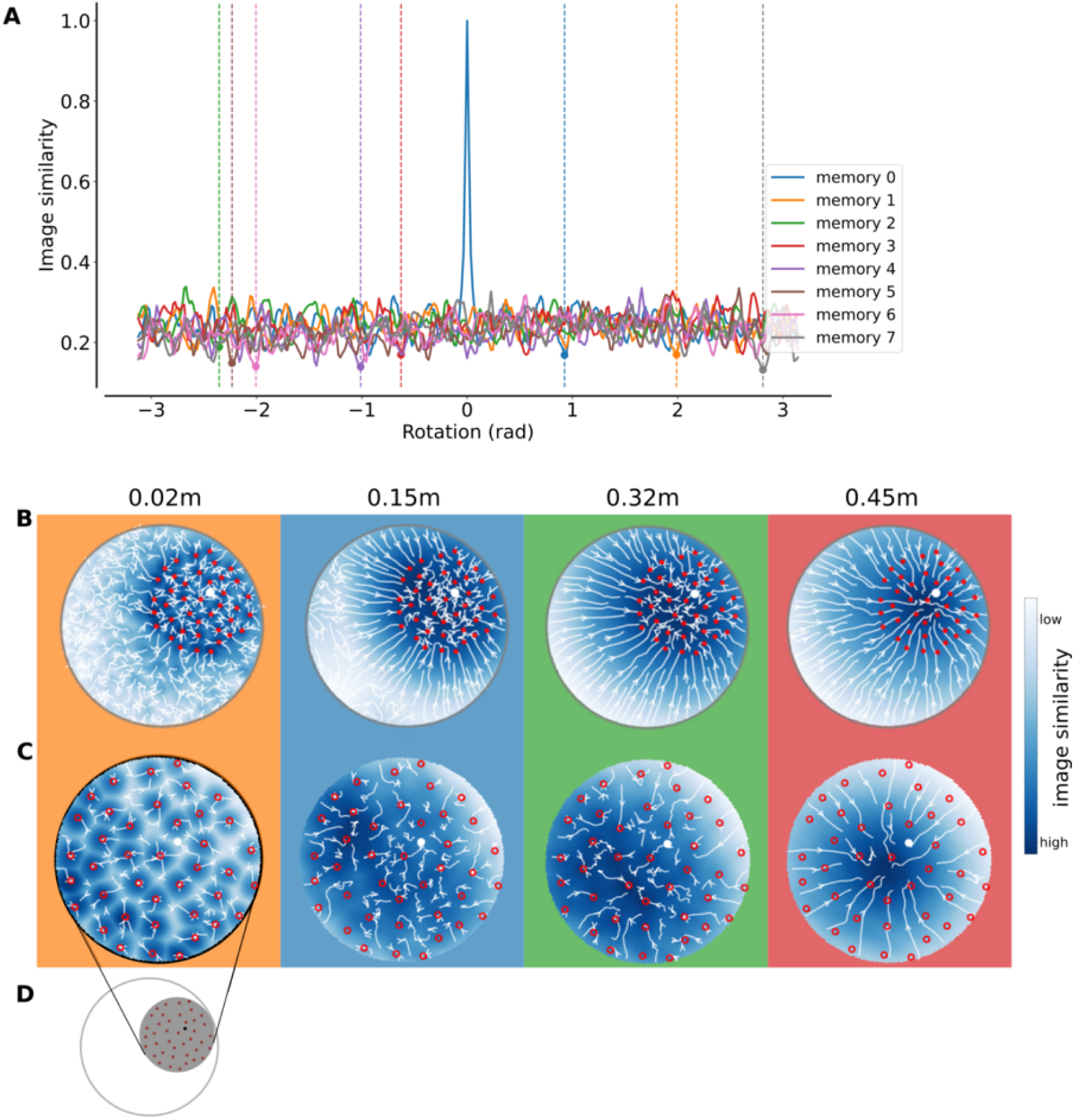
Contrast-weighted nearness based homing model at four altitudes (0.02 m, 0.15 m, 0.32 m, and 0.45 m) with bird’s eye view snapshots taken inside the dense environment (distance = 0.1 m, height = 0.45 m). **A:** We applied the model to a list of images, these images are examples of the eight memorised snapshots which are aligned in the nest direction and taken around the nest. For the rotational image difference function, we used memory 0 as a reference image, and compared the seven others by rotating them against memory 0. We observe that the image similarity as based on [14] is the maximum for the memorised image at a null rotation, as expected. If the other images are not two far from the nest, we may see other local maxima for each of these image, where the local maxima are shifted according the nest bearing. **B-D:** Heatmaps of full environment (**B**) and only the dense area (**C**, as shown in **D**) of the image similarity. The colour indicates the image similarity between the view at the position in the arena and the snapshots taken around the nest. The model leads the simulated bees with bird’s eye view snapshots inside the dense environment only to the center of the dense environment (altitude of 0.45 m) but not to the nest.

**Figure 15.**
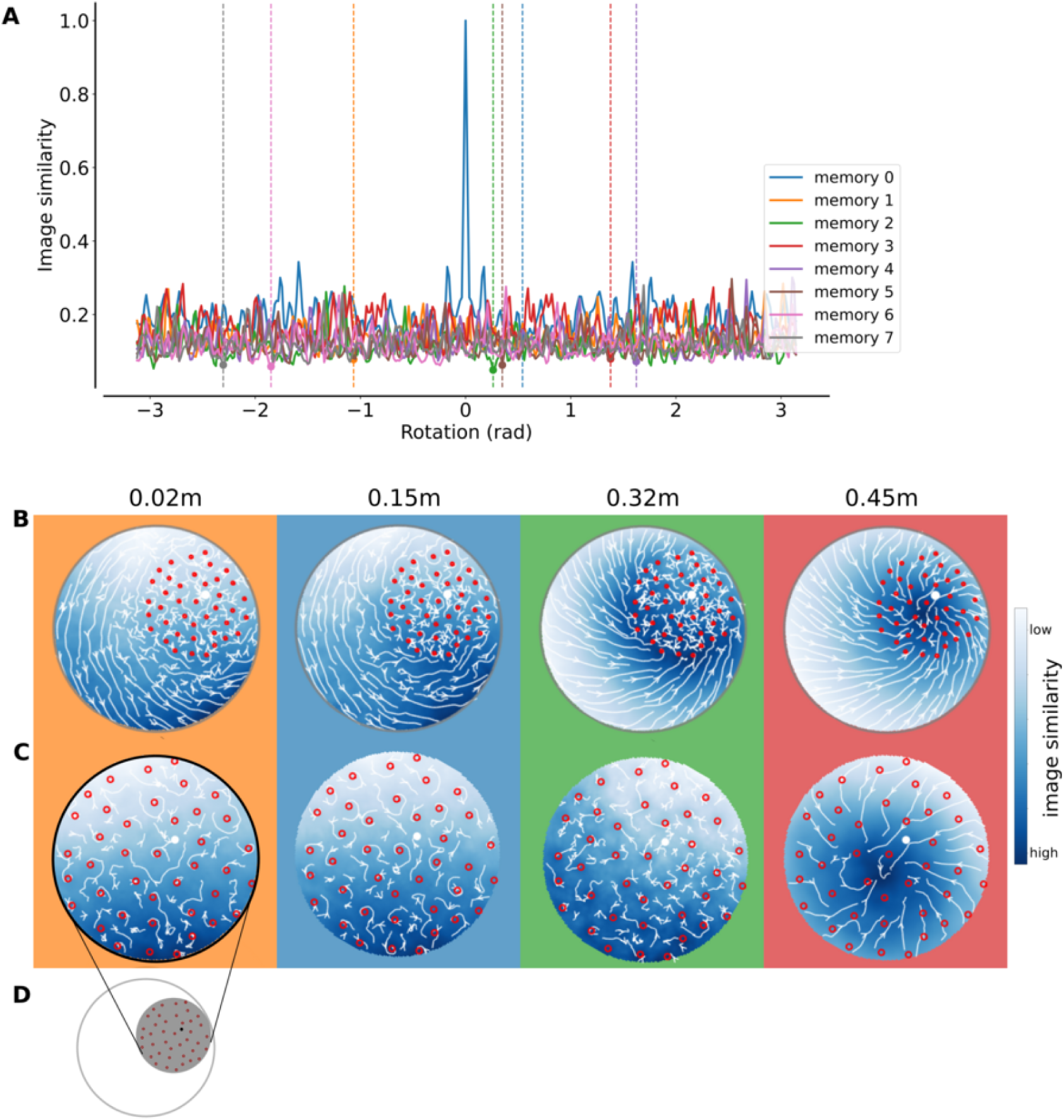
Contrast-weighted nearness based homing model at four altitudes (0.02 m, 0.15 m, 0.32 m, and 0.45 m) with ground view snapshots taken inside the dense environment (distance = 0.1 m, height = 0.02 m). The colour indicates the image difference between the view at the position in the arena and the snapshots taken around the nest. **A:** We applied the model to a list of images, these images are examples of the eight memorised snapshots which are aligned in the nest direction and taken around the nest. For the rotational image difference function, we used memory 0 as a reference image, and compared the seven others by rotating them against memory 0. We observe that the image similarity as based on [14] is the maximum for the memorised image at a null rotation, as expected. If the other images are not two far from the nest, we may see other local maxima for each of these images, where the local maxima are shifted according the nest bearing. **B-D:** Heatmaps of full environment (**B**) and only the dense area (**C**, as shown in **D**) of the image similarity. The colour indicates the image similarity between the view at the position in the arena and the snapshots taken around the nest. The model leads the simulated bees with ground view snapshots inside the dense environment only to the center of the dense environment (altitude of 0.45 m) but not to the nest.

**Figure 16.**
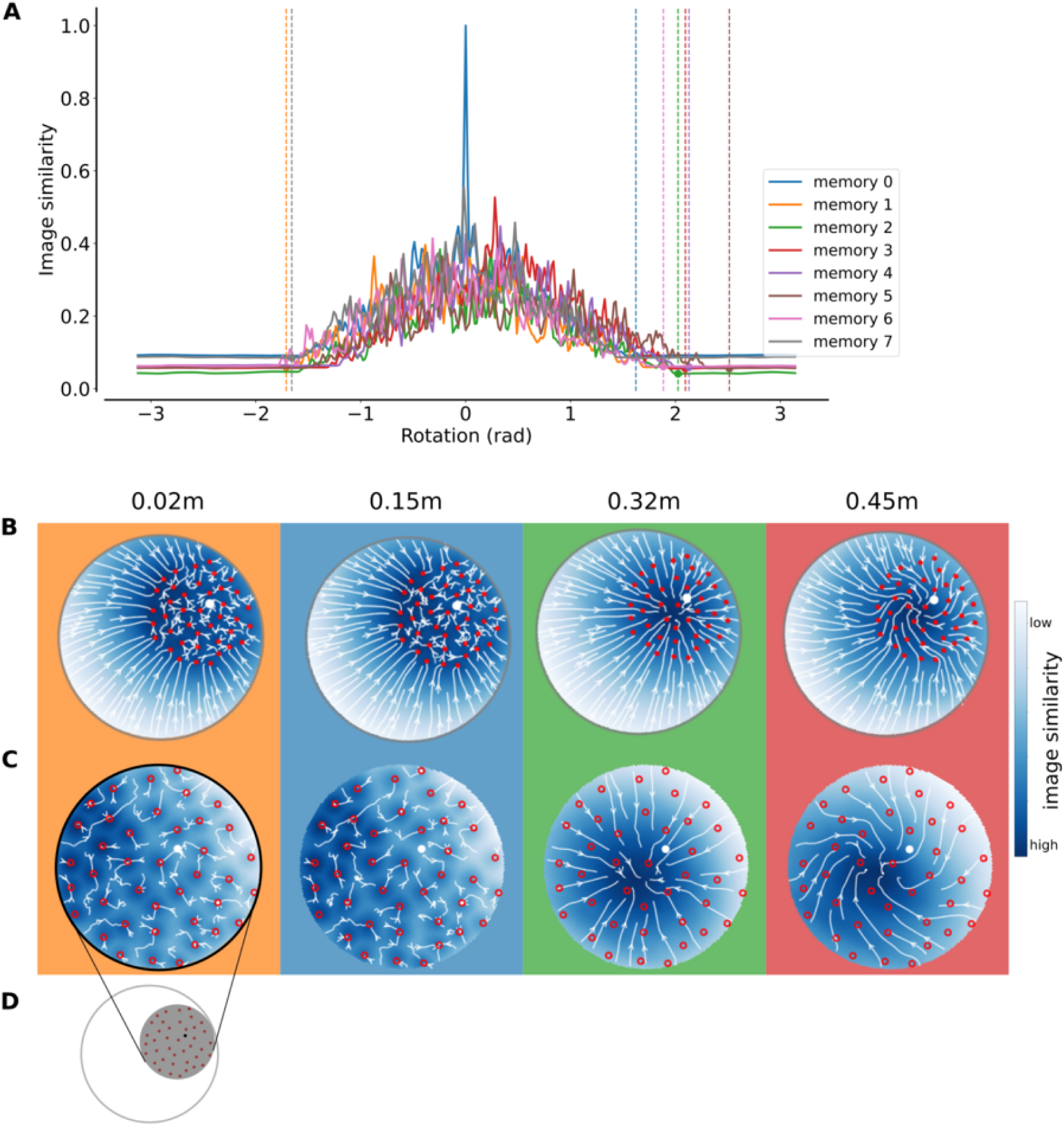
Contrast-weighted nearness based homing model at four altitudes (0.02 m, 0.15 m, 0.32 m, and 0.45 m) with ground view snapshots taken outside the dense environment (distance = 0.55 m, height = 0.02 m). The colour indicates the image difference between the view at the position in the arena and the snapshots taken around the nest. **A:** We applied the model to a list of images, these images are examples of the eight memorised snapshots which are aligned in the nest direction and taken around the nest. For the rotational image difference function, we used memory 0 as a reference image, and compared the seven others by rotating them against memory 0. We observe that the image similarity as based on [14] is the maximum for the memorised image at a null rotation, as expected. If the other images are not two far from the nest, we may see other local maxima for each of these images, where the local maxima are shifted according the nest bearing. **B-D:** Heatmaps of full environment (**B**) and only the dense area (**C**, as shown in **D**) of the image similarity. The colour indicates the image similarity between the view at the position in the arena and the snapshots taken around the nest. The model leads the simulated bees with ground view snapshots outside the dense environment only to the center of the dense environment (altitude of 0.32 m) or slightly shifted away from center (altitude of 0.45 m) but not to the nest.

**Figure 17.**
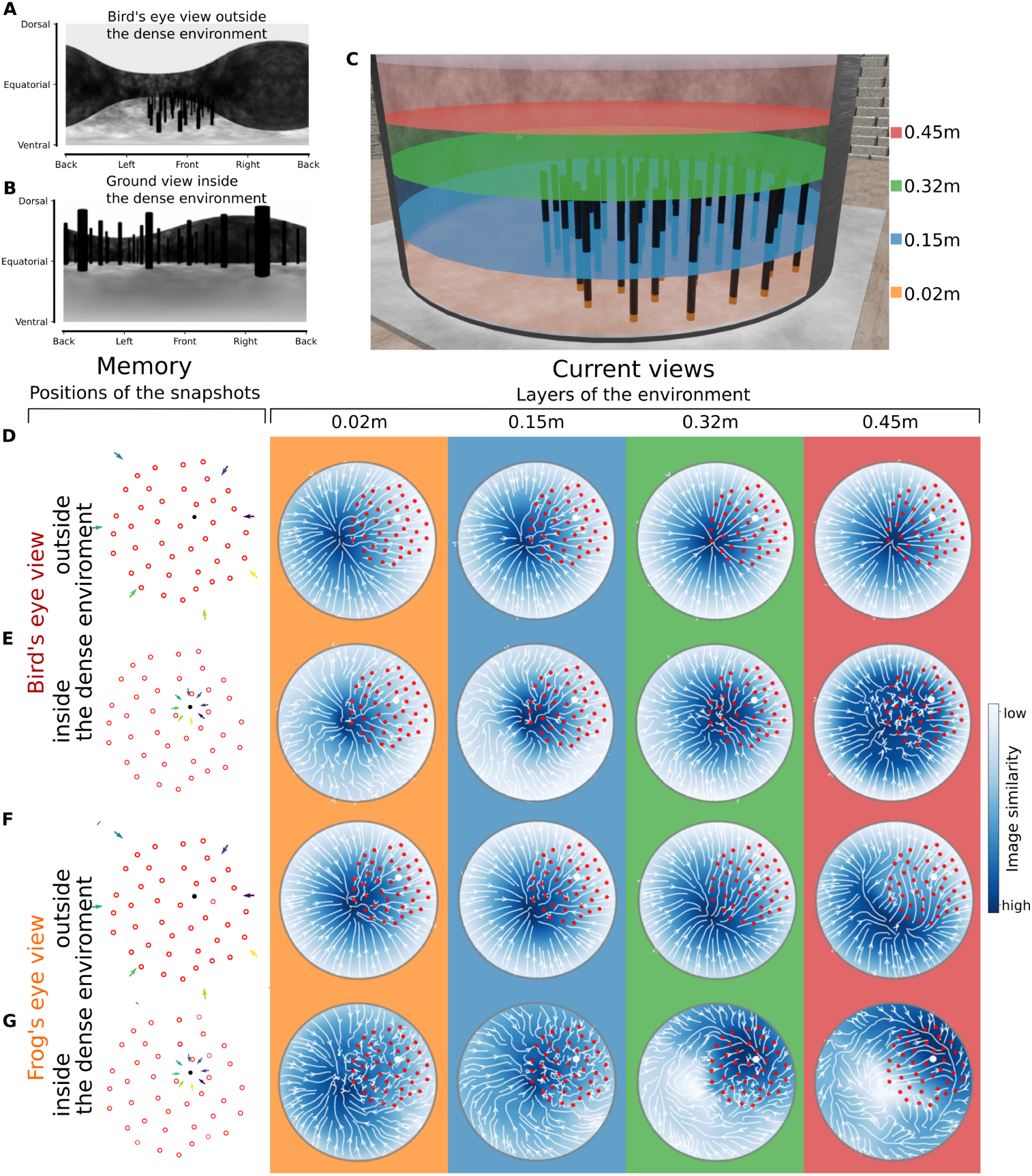
Brightness based homing model with the simulated wall of the cylinder at four altitudes (0.02 m, 0.15 m, 0.32 m, and 0.45 m) with bird’s eye view (**D&E**) and ground view (**F&G**) snapshots taken outside (**D&F**) and inside (**E&G**) the dense environment. **A&B:** Examples of panoramic snapshots in A from the bird’s eye view outside the dense environment and in B ground views inside the dense environment. The axes of the panoramic images refer to the azimuthal directions (x-axis) and to the elevational directions from the simulated bee’s point of view (y-axis, dorsal meaning upwards, ventral downwards, equatorial towards the horizon). **C:** Rendered layers of the environment for a comparison of the current view of the simulated bee. The layers are at 0.02 m(orange), 0.15 m (blue), 0.32 m(green) and 0.45 m (red) heights. **D-G:** The first column shows were the snapshots were taken in relation to the nest position (nest position in black, objects in red and snapshot positions indicated by coloured arrows). The other four columns show the comparison of memorised snapshot the four layers of the environment (as shown in **C**). The heatmaps show the image similarity between the current view at the position in the arena and the memorised snapshots taken around the nest (blue = very similar, white = very different). Additionally, white lines and arrows present the vector field from which the homing potential is derived. Red circles indicate the positions of the objects and the white dot indicates the nest position. The background colour of each column indicates the height of the current views that the snapshots are compared to.

**Figure 18.**
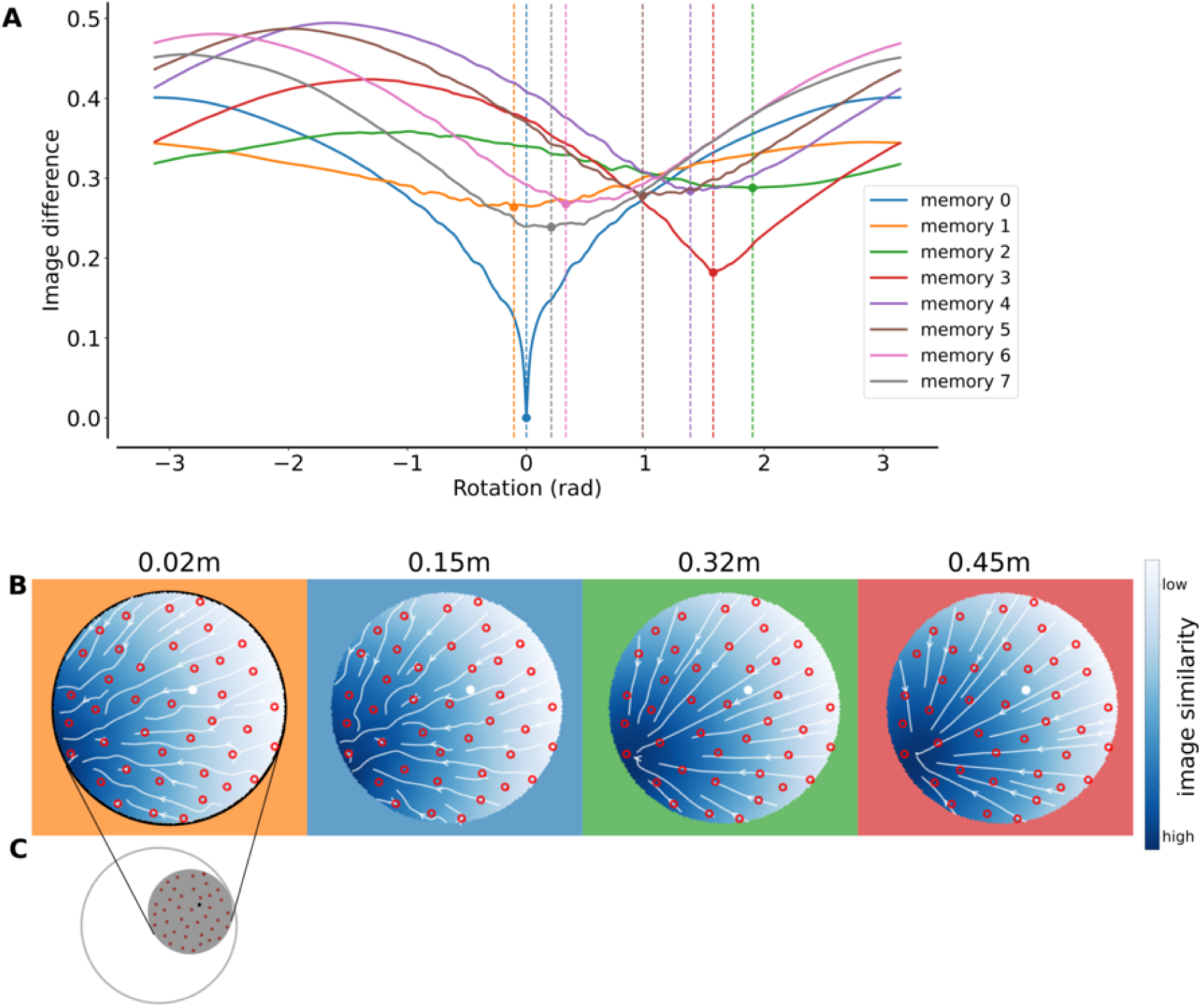
Brightness-based homing model with the simulated wall of the cylinder at four altitudes (0.02 m, 0.15 m, 0.32 m, and 0.45 m) with bird’s eye view snapshots taken outside the dense environment (distance = 0.55 m, height = 0.45 m). **A:** We applied the model to a list of images, these images are examples of the eight memorised snapshots which are aligned in the nest direction and taken around the nest. For the rotational image difference function, we used memory 0 as a reference image, and compared the seven others by rotating them against memory 0. We observe that the image difference function is minimum for the memorised image at a null rotation, as expected. If the other images are not two far from the nest, we may see other local minima for each of these images, where the local minima are shifted according the nest bearing. **B-C:** Heatmaps of the dense area (**B**, as shown in **C**) of the image similarity. The colour indicates the image difference between the view at the position in the arena and the snapshots taken around the nest.

**Figure 19.**
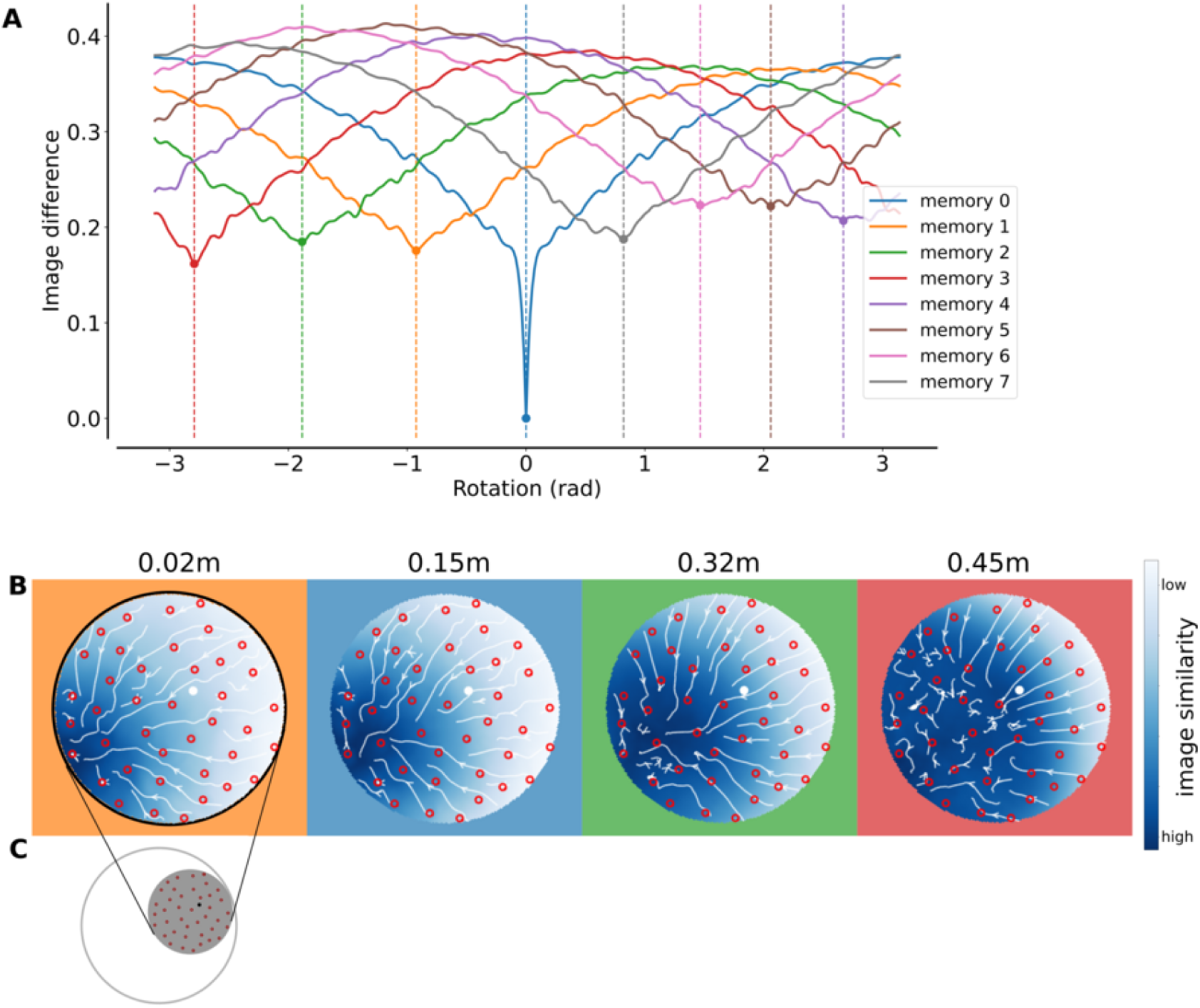
Brightness-based homing model with the simulated wall of the cylinder at four altitudes (0.02 m, 0.15 m, 0.32 m, and 0.45 m) with bird’s eye view snapshots taken inside the dense environment (distance = 0.1 m, height = 0.45 m). The colour indicates the image difference between the view at the position in the arena and the snapshots taken around the nest. **A:** We applied the model to a list of images, these images are examples of the eight memorised snapshots which are aligned in the nest direction and taken around the nest. For the rotational image difference function, we used memory 0 as a reference image, and compared the seven others by rotating them against memory 0. We observe that the image difference function is minimum for the memorised image at a null rotation, as expected. If the other images are not two far from the nest, we may see other local minima for each of these images, where the local minima are shifted according the nest bearing. **B-C:** Heatmaps of the dense area (**B**, as shown in **C**) of the image similarity. The colour indicates the image difference between the view at the position in the arena and the snapshots taken around the nest.

**Figure 20.**
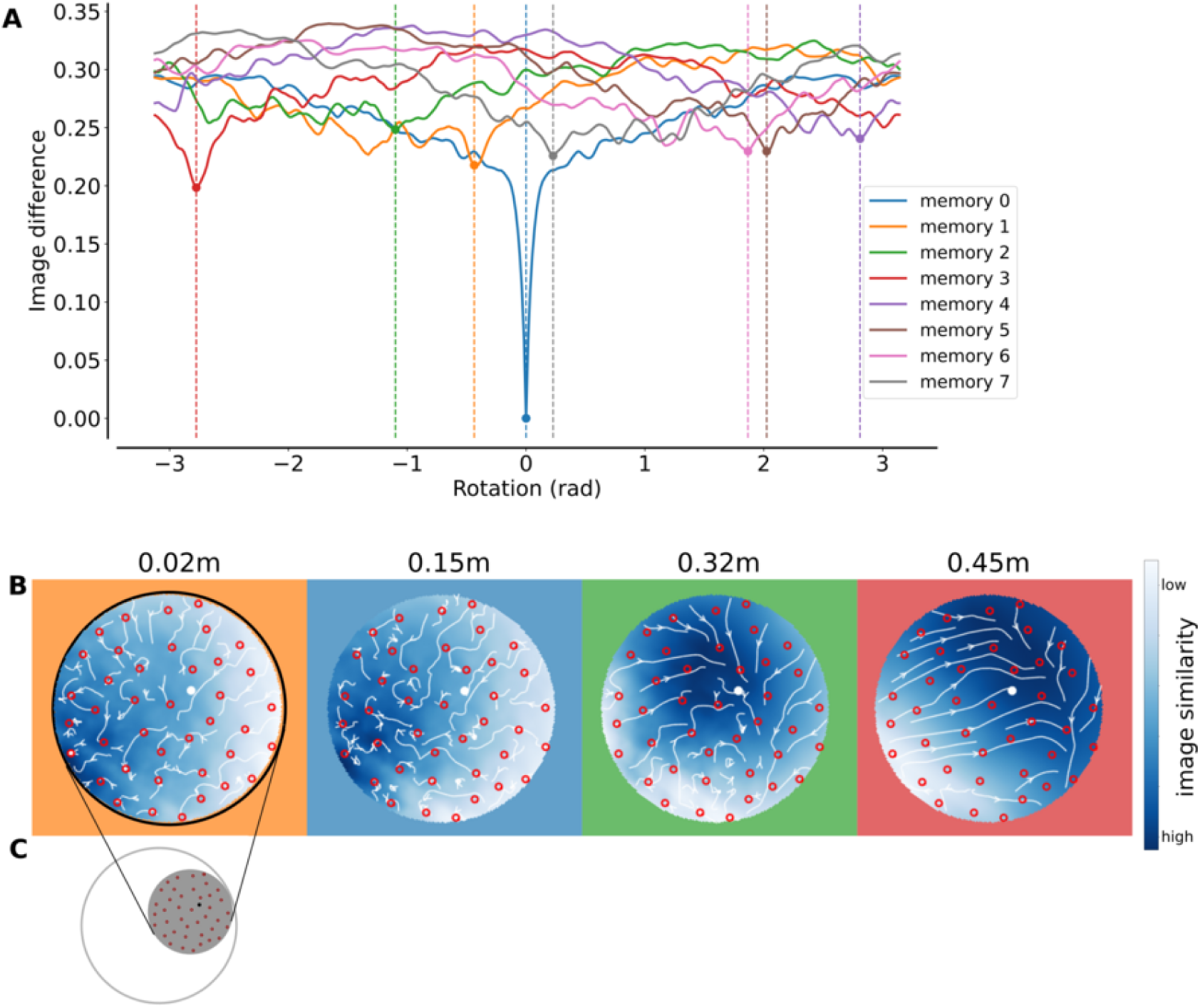
Brightness-based homing model with the simulated wall of the cylinder at four altitudes (0.02 m, 0.15 m, 0.32 m, and 0.45 m) with ground view snapshots taken inside the dense environment (distance = 0.1 m, height = 0.02 m). The colour indicates the image difference between the view at the position in the arena and the snapshots taken around the nest. **A:** We applied the model to a list of images, these images are examples of the eight memorised snapshots which are aligned in the nest direction and taken around the nest. For the rotational image difference function, we used memory 0 as a reference image, and compared the seven others by rotating them against memory 0. We observe that the image difference function is minimum for the memorised image at a null rotation, as expected. If the other images are not two far from the nest, we may see other local minima for each of these images, where the local minima are shifted according the nest bearing. **B-C:** Heatmaps of the dense area (**B**, as shown in **C**) of the image similarity. The colour indicates the image difference between the view at the position in the arena and the snapshots taken around the nest.

**Figure 21.**
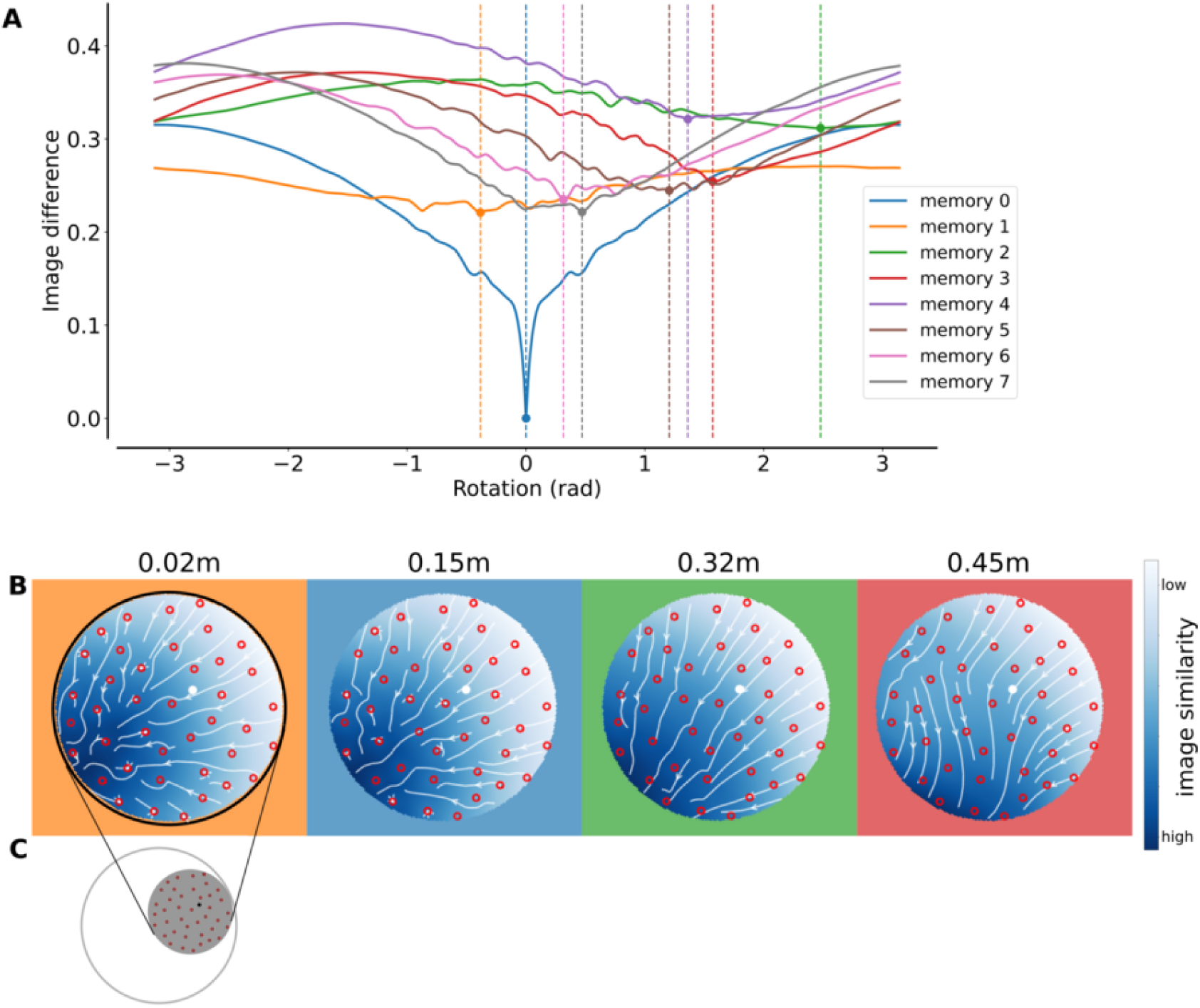
Brightness-based homing model with the simulated wall of the cylinder at four altitudes (0.02 m, 0.15 m, 0.32 m, and 0.45 m) with ground view snapshots taken outside the dense environment (distance = 0.55 m, height = 0.02 m). The colour indicates the image difference between the view at the position in the arena and the snapshots taken around the nest. **A:** We applied the model to a list of images, these images are examples of the eight memorised snapshots which are aligned in the nest direction and taken around the nest. For the rotational image difference function, we used memory 0 as a reference image, and compared the seven others by rotating them against memory 0. We observe that the image difference function is minimum for the memorised image at a null rotation, as expected. If the other images are not two far from the nest, we may see other local minima for each of these image, where the local minima are shifted according the nest bearing. **B-C:** Heatmaps of the dense area (**B**, as shown in **C**) of the image similarity. The colour indicates the image difference between the view at the position in the arena and the snapshots taken around the nest.

**Figure 22.**
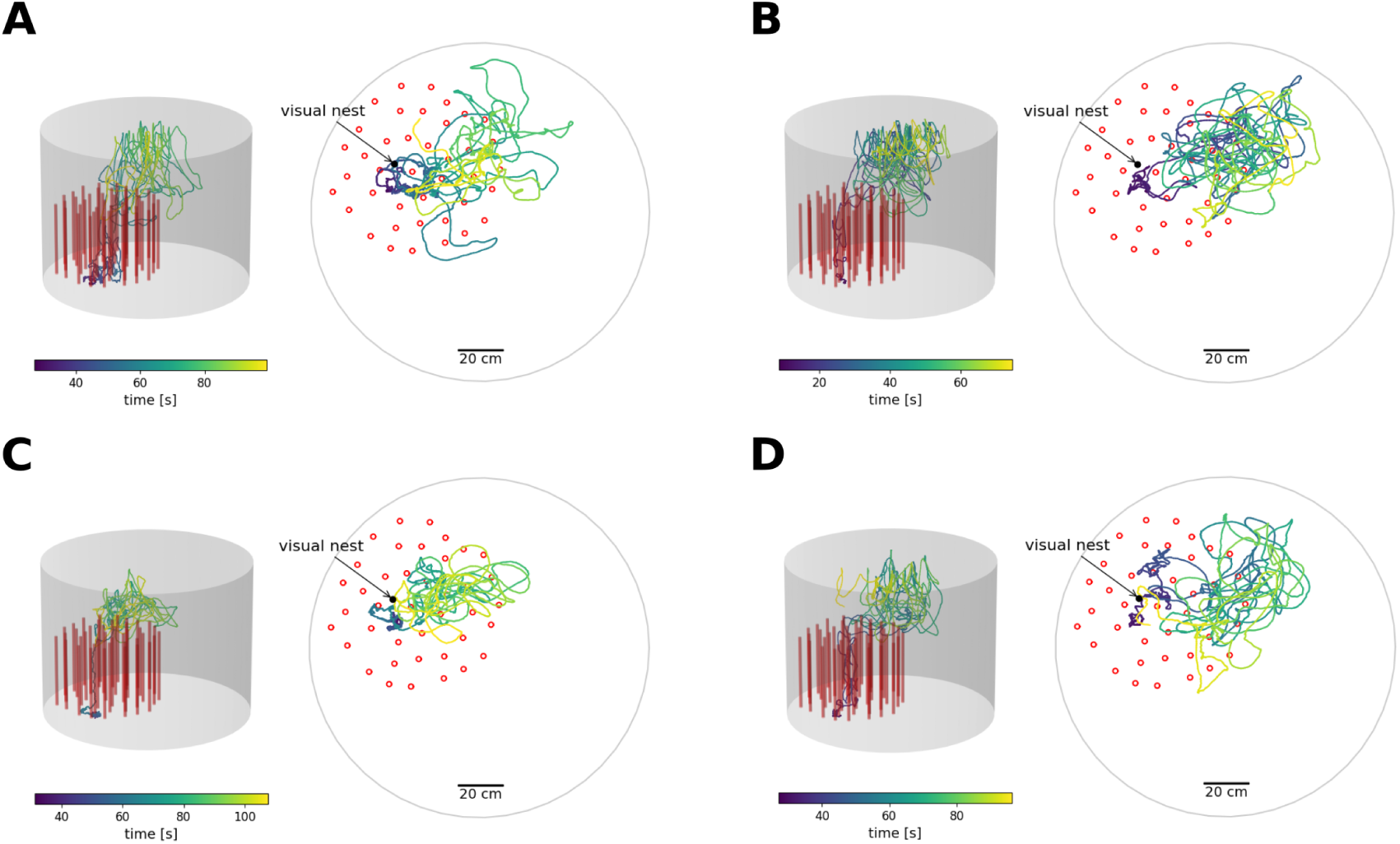
Four exemplary flight trajectories of the first outbound flight of bees in 3D (left subplot) and a top view in 2D (right subplot). The colour indicates the time, blue the start of the and red the end of the flight. The objects are depicted by red cylinders in the 3D plot and as red circles in the 2D plot. The black dots in the 2D plot shows the nest position within the dense environment.

**Figure 23.**
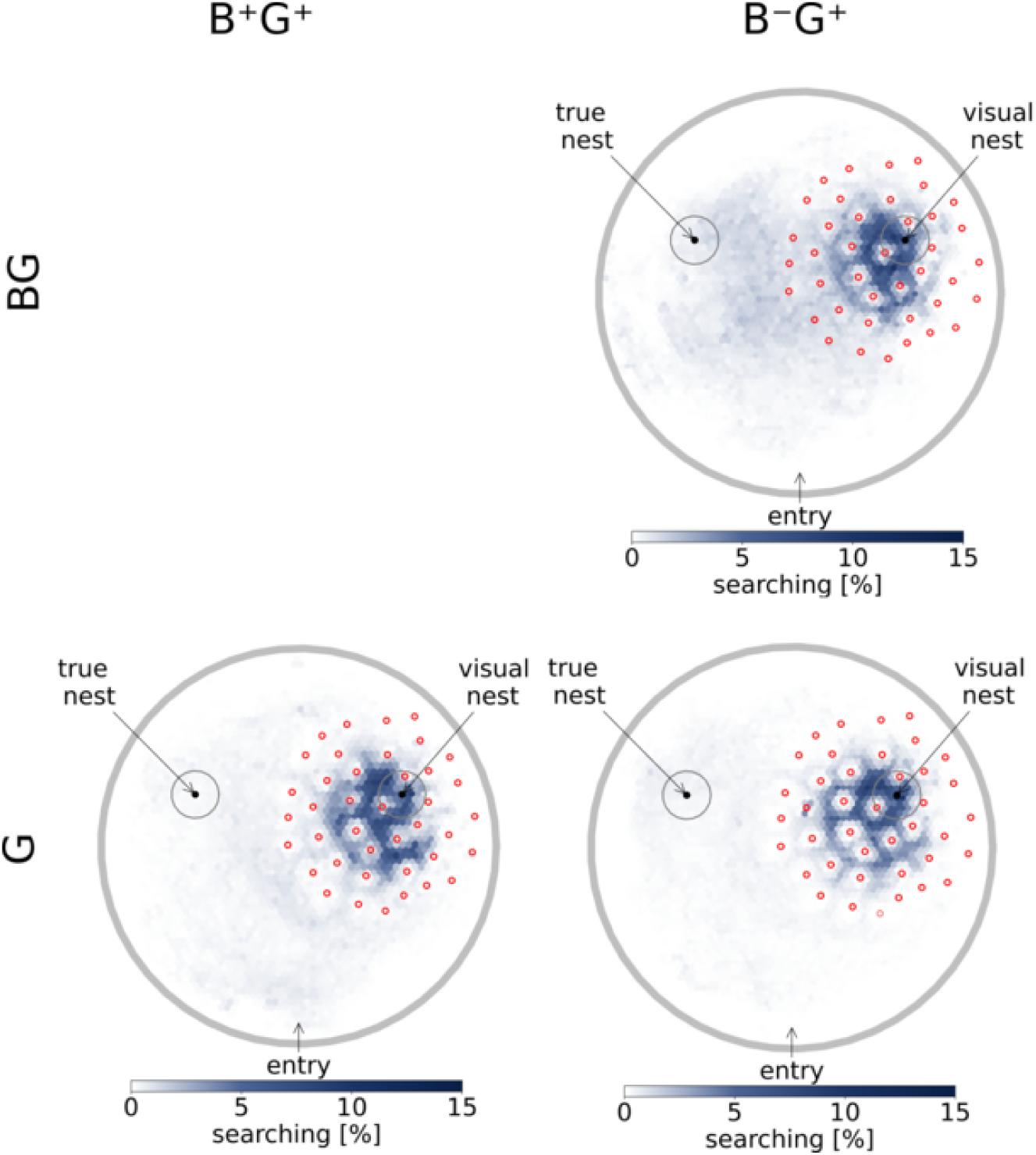
Search distributions for the group B^+^G^+^ (left column) in the condition G (condition BG is shown in the main results in Fig.2D) and for group B*^−^*G^+^ (right column) in the tests BG and G (N = 26 for each plot). The bees of group B^+^G^+^ in the condition G searched for the nest inside the dense environment and spent more time at the visual nest (in the dense environment) than at the true nest (as during training). The bees of group B*^−^*G^+^ in the condition G and BG searched for the nest within the dense environment and spent more time at the visual nest than at the true nest.

**Figure 24.**
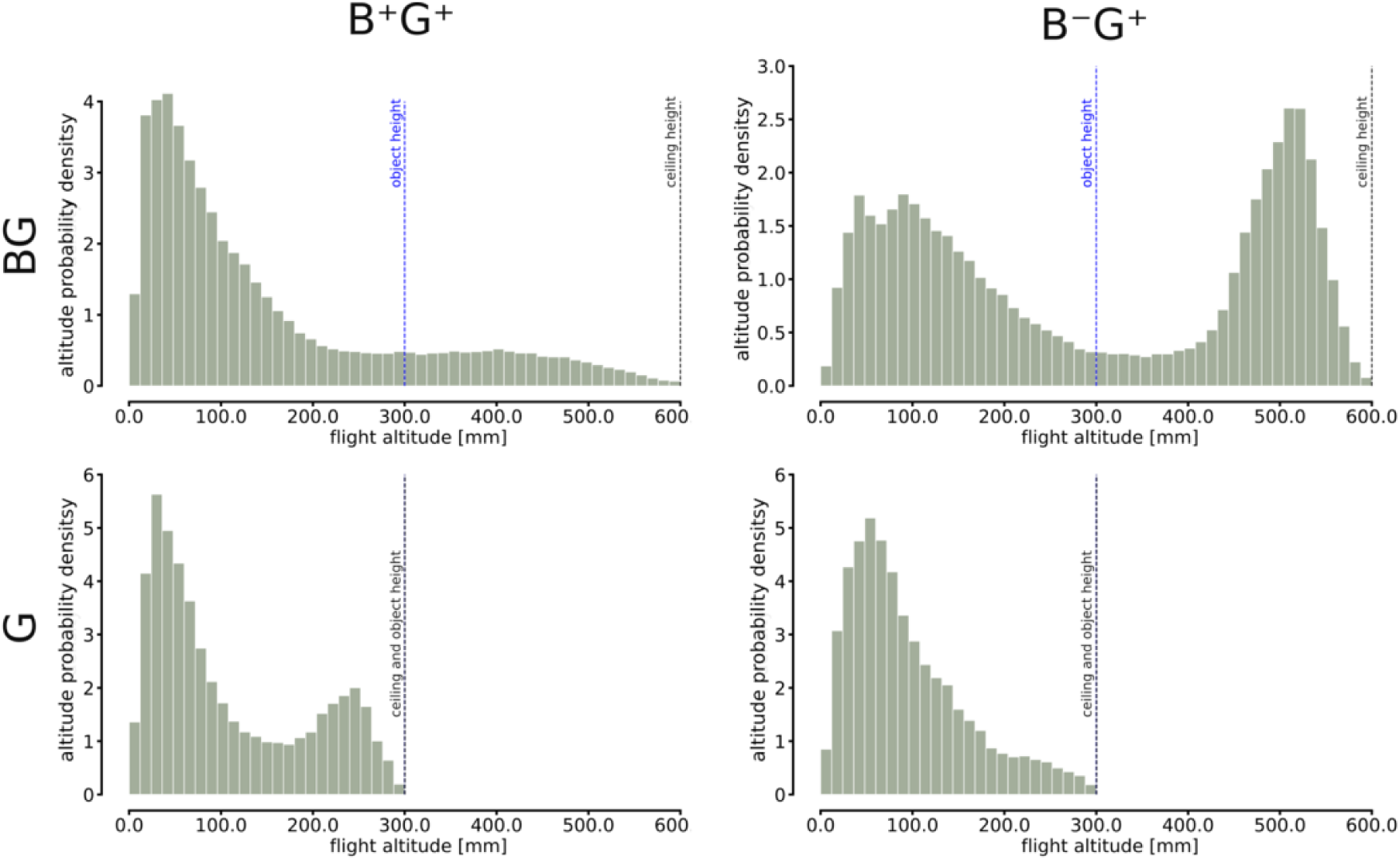
Flight altitude distributions for the tests BG and G for the groups B^+^G^+^ and B*^−^*G^+^ (N = 26 for each plot).

**Figure 25.**
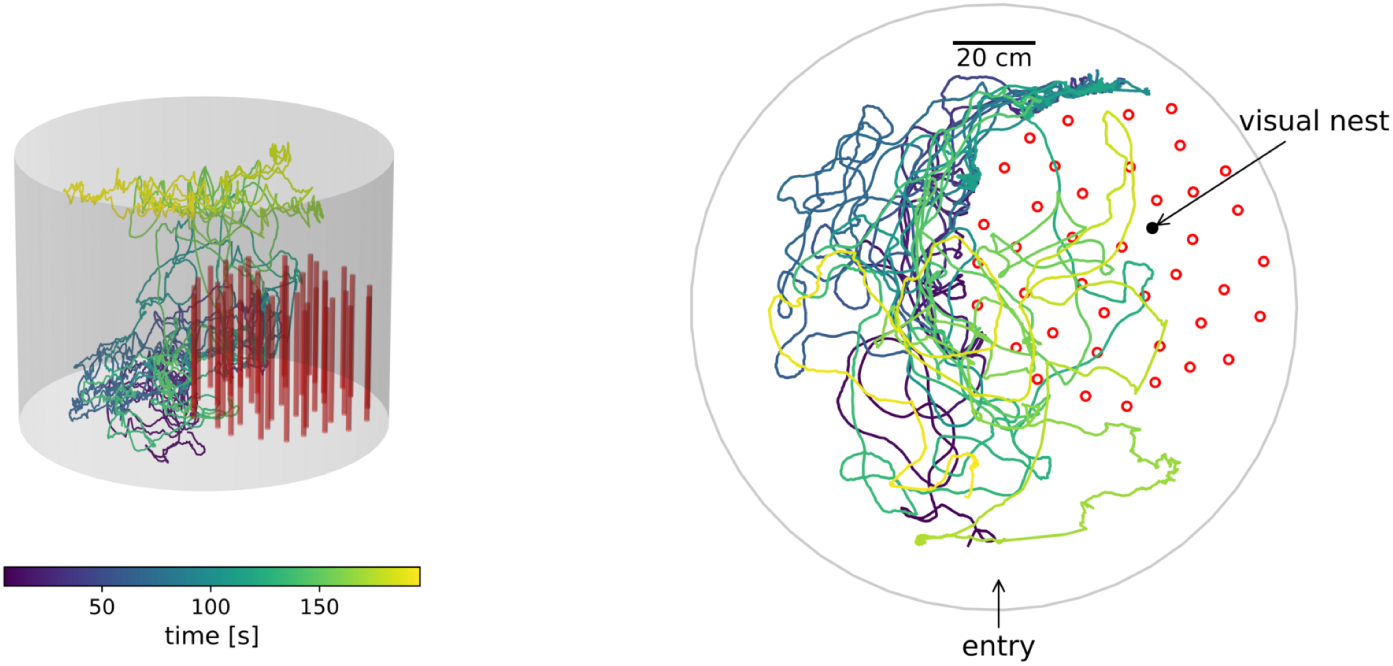
Exemplary flight trajectories in 3D (left column) and a top view in 2D (right column) from the group B^+^G^+^ in the B test. The colour indicates the time, blue the start of the and red the end of the flight. The objects are depicted by red cylinders in the 3D plot and as red circles in the 2D plot. The black dots in the 2D plot shows the nest position within the dense environment. The bee is trying to enter the covered dense environment from the side but, eventually, it is increasing its altitude and it flies above the dense environment. However, the bee is not searching for the visual nest above the covered dense environment.

We used the method presented in Doussot et al. [15] to calculate the image differences according to the brightness and the contrast-weighted-nearness method, each with eight snapshots. The image difference was calculated either by the brightness method relying on the brightness value of each pixel in one panoramic image or the contrast-weighted nearness method (see SI Methods) relying on the contrast calculated by the Michelson contrast (ratio of the luminance-amplitude (I*_max_ −* I*_min_*)) and the luminance-background (I*_max_* +I*_min_*)) weighted by the inverse of the distance [14]. We wanted to predict the simulated bee’s probable endpoint by analyzing vector fields from different homing models. Convergence points in these vector fields were determined using the Helmholtz-Hodge Decomposition, focusing on the curl-free component, representing the potential ϕ ([15]). After applying the Helmholtz-Hodge Decomposition to each vector field, we scaled the resulting potential between 0 and 1 for all homing models. With these potentials, we were able to plot heatmaps accordingly to estimate the areas in the arena where view-based agents/ simulated bees would most likely steer towards. These heatmaps were compared to the bees’ search distributions in the arena.

### Animal handling

We used four *B. terrestris* hives provided by Koppert B.V., The Netherlands, that were ordered sequentially to test one colony at a time. The bee hives arrived in small boxes and were transferred to acrylic nest boxes under red light (non-visible to bees [16]) to 30 *×* 30*×* 30 cm. The nest box was placed on the floor and connected to an experimental arena. We covered the nest box with a black cloth to mimic the natural lighting of underground habitats of *B. terrestris* [20]. The bee colonies were provided with pollen balls *ad libitum* directly within the nest boxes. The pollen balls were made of 50 mL ground, commercial pollen collected by honeybees (W. Seip, Germany), and 10 mL water. The bees reached a foraging chamber via the experimental arena containing gravity feeders. These are small bottles with a plate at the bottom so that the bees can access the sugar solution through small slots in the plate. The feeders were filled with a sweet aqueous solution (30% saccharose, 70% water in volume). Lighting from above was provided in a 12 h/12 h cycle and the temperature was kept constantly at 20*^◦^*. Throughout the experiments, foraging bees were individually marked using coloured, numbered plastic tags glued with melted resin on their thorax.

### Experimental arena

The experimental arena was a cylinder with a diameter of 1.5 m and a height of 80 cm as in [15]. A red and white pattern, perceived black and white by bumblebees [16], with a 1/f spatial frequency distribution (a distribution observed in nature [40]) covered the wall and floor of the cylindrical arena; the bees were provided with enough contrast to allow them to use optical flow. Red light came from 18 neon tubes (36 W Osram + Triconic) filtered by a red acrylic plate (Antiflex ac red 1600 ttv). Bees did not see these lights and perceived the red-white pattern as black and white. An adjustable transparent ceiling prevented the bees from exiting the arena. It allowed lighting from 8 neon tubes (52 W Osram + Triconic) and 8 LEDs (5 W GreenLED) as in [15], and recording from five high-speed cameras with different viewing angles. As in many previous indoor experiments about bee navigation, UV light was not provided within the arena [30, 15, 19]. A foraging bee exiting the hive crossed three small acrylic boxes (inner dimension of 8 *×* 8*×* 8 cm) with closable exits to select the bee to be tested. The bee then walked through a plastic tube with 2.5 cm in diameter and entered the cylindrical arena. This nest exit was surrounded by visual objects. The foraging chamber was reached through a hole at a height of 28 cm above the floor in the arena wall. After foraging the bees exited the foraging chamber and entered the flight arena through a hole in the arena wall at a height of 28 cm above the floor. The nest entrance within the arena was surrounded by visual objects. When it found this nest entrance, the bee walked through a plastic tube with 2.5 cm in diameter and reached the hive by crossing three small acrylic boxes (inner dimension of 8 *×* 8*×* 8 cm) with closable exits which were used during the experiments to block bees from entering the arena. The objects surrounding the nest exit in the arena consisted of 40 randomly placed red cylinders (2 cm in diameter and 30 cm in height), creating an artificial meadow. The size of the dense object constellation was 80 cm in diameter to allow its displacement to a different location in the flight arena. We considered that the object constellation was dense enough to pose a challenge in finding the nest within the cylinder constellations, as we could show in the snapshot-model comparison. For example, if only three cylinders were used, the bee may only search there because these would be the only conspicuous landmarks. Additionally, the configuration had to be sufficiently sparse for the bee to fly through [38]. We used the object density and object distances of Gonsek et al. [19] as a reference to find a randomly distributed object configuration. Red-lighting from below was used only during recordings. The cylindrical arena had a door, allowing the experimenter to change the objects within the arena.

### Experimental design

We tested two groups of bees in three tests trained with the dense object constellation (dense environment) surrounding the nest entrance. The two groups differed in the training condition. For the group B^+^G^+^, the flight altitude was unconstrained, and they could experience bird’s eye views above the objects. The second group, B*^−^*G^+^, were restricted during training to a maximum flight altitude of the height of the objects, so the ground views.

We tested 26 bees per group (4 colonies were used, 2 per group, from each colony 13 bees are included in the analysis resulting in a total of 52 individuals). In both groups, the foraging bees travelled between their nest and the foraging chamber. The return flight of each bee was recorded in all tests of a given experiment. Between individual tests, the bees were allowed to forage *ad libitum*. A dense object constellation surrounded the nest, and the area around the nest positions was cleaned with 70% ethanol between the tests to avoid chemical markings.

To test the behaviour of the bees, we locked up to six individually marked bees in the foraging chamber at a time. Each bee participated either in group B^+^G^+^ (high ceiling during training, resulting in available ground and bird’s eye views) or B*^−^*G^+^ (low ceiling during training, resulting in only available ground views) and was tested once in all tests of the respective experiment. The order of the tests was pseudo-random only that in the group B*^−^*G^+^ the test BG was always tested last to not let the bees experience bird’s eye views before testing the B test. Before the tests, the cylindrical arena was emptied of bees, the spatial arrangement of objects was shifted, and the nest entrance closed. One bee at a time was allowed to search for its home for three minutes after take-off. After this time, the spatial arrangement of the ceiling height and the dense object constellation was placed back in the training condition, and the nest entrance opened. The bees had up to two minutes to take off. Otherwise, they were captured and released close to the nest. Between tests, the bees could fly *ad libitum* between the nest and foraging chamber under the training conditions.

For the tests, the dense environment was placed at a different location than during the training condition, and thus, it did not surround the true nest entrance leading to the hive. The dense environment indicated the location of a visual nest entrance. The true nest entrance and the visual nest entrance were covered by a piece of paper with the same texture as the arena floor so that they were not discernible by the bees.

For the test BF, no other constraint was added to the general tests to test if the bees associated their nest entrance with the dense environment. The G test consisted of a transparent wall and ceiling on top of the objects, which prevented the bees from entering the dense object constellation. To return to the nest location in the dense object constellation, they were only able to pinpoint the position from above if they were using only bird’s eye views. In the G test, the flight altitude was constrained to the height of the dense environment so that bees could no longer experience bird’s eye views from above the dense environment. Thus, the bees had to use the ground view to return to their nest.

### Flight trajectories

Bee trajectories were recorded at 62.5 Hz (16 ms between two consecutive frames) with five synchronised Basler cameras (Basler acA 2040um-NIR) with different viewing angles (similar to [15, 35]). One camera was placed on top of the middle of the arena to track the bumblebees’ movements in the plane, and the other four cameras were distributed around the arena to record the position of the bees from different angles to minimise the error during triangulation of the bee’s position in 3D. Before the bees entered the setup, the recording had already started, and the first 60 frames were used to calculate a bee-less image of the arena (background image). During the rest of the recording, only crops, i.e. image sections (40×40 pixels), containing large differences between the background image and the current image (i.e. potentially containing the bee) were saved to the hard drive together with the position of the crop in the image. The recording scripts were written in C++. The image crops were analysed by a custom-written neural network to classify the crops in bees or not a bee. When non-biological speed (speed above 4 m/s [20]) or implausible positions (outside the arena) were observed, the crops neighbouring these time points were manually reviewed.

The trajectories were analysed in Python (version 3.8.17), notably with OpenCV. A detailed list of packages used is published in the data publication. The time course of the positions of the bees in 3D within the arena is shown for a selection of flights (Fig. 2 and S13). For each of the tests, a distribution of presence in the flight arena was computed by using hexagonal binning of the 2D positions to show the search areas of the bees qualitatively. Bees that collected food in the foraging chamber returned home and searched for their nest entrance. Even when objects are displaced to a novel location [27, 28, 39, 29], replaced by smaller, differently coloured, or camouflaged objects [27, 28, 14], or moved to create visual conflict between several visual features [15], bees search for their nest entrance associated to such objects. Therefore, we assumed that bees entering the arena after visiting a foraging chamber will search for their nest location. The returning bees could spend an equal amount of time in any location in the arena or concentrate their search around the physical position of their nest or around visual objects associated with the nest entrance [15]. In our experiments, the nest entrance was surrounded by cylindrical objects. During the tests, when the objects were shifted to a novel location, a bee guided by the objects might have searched for its nest around the objects. In contrast, they might not have used the objects to navigate and might have searched for their nest at the original location. We, therefore, quantified the time spent around two locations: at the true nest and the nest according to the objects (‘visual nest’). Additionally, the positions of the bees crossing the boundaries of the densely distributed objects for the first time at the side or at the top of the dense environment were visualised a circular histogram (entries from the top) and a scatter plot (entries from the side). These were used to describe where the bees entered the dense area. As the dense environment was shifted to another position in the arena, we can exclude the use of compass, odour or magnetic cues.

### Statistical analysis for hypotheses testing

Hypotheses about the time spent in one area compared to another were tested using the dependent t-test for paired samples. Hypotheses involving multiple comparisons were tested using a Bonferroni correction for the significance level. As long as a hypothesis concerned only two areas, no adjustment was made to the significance level. With a sample size of 26 bees, we were able to detect a time difference spent in two areas (e.g. fictive and true nest locations) of 0.25 s assuming a standard deviation of 0.38 s (estimated from [15]) with a power of 80% at a significance level of 0.05. The analysis was performed with Python using the Scipy library for statistical analyses.

## Acknowledgments

We would like to thank Vedant Dixit, Maximilian Stahlsmeier, Pia Hippel and Helene Schnellenberg for their help during the data collection. Additionally, we would like to thank Sina Mews for helpful discussions on statistical models. This project was supported by the Deutsche Forschungsgemeinschaft (DFG, German Research Foundation) and the Agence Nationale de la Recherche (ANR, French National Research Agency). We also acknowledge support for the publication costs by the Open Access Publication Fund of Bielefeld University.

## Supporting Information

### Methods

#### Rotational Image Difference Functions

For each position (*x, y*) in the arena an equi-rectangular panoramic image (360deg along the azimuth, 180 deg along the elevation) was acquired. To determine the most familiar direction, each snapshot in the arena were compared to memorised snapshot. This comparison was based on the minimum rotational image difference function (RIDF, Eq. 1) for the brightness model. The minimum RIDF is the minimum root mean squared image difference d*_x,y_* between two views (the current view *I_x,y_* and the view at the nest *I_N_*) for different azimuthal viewing directions α weighted by *w*(*v*). *w*(*v*) is a sine wave along the y-axis counterbalancing the oversampling of the poles at the transformation of the 3D sphere mimicking the bee’s eye to 2D equirectangular images by giving values of 1 at the equator and 0 at the poles (Eq. 2). In the equation below, (*u, v*) corresponds to the viewing direction in the azimuthal direction u, and direction along the elevation v. The images resolution were (*N_u_, N_v_*) = (360, 180) pixels.

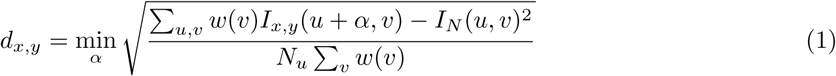

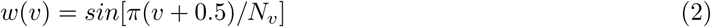

The RIDF *d_si,x,y_* is calculated for each snapshot *s_i_* in *S* = *s*_0_, *s*_1_, …, *s_n_* around the nest location and the heading direction *h_si,x,y_* at each grid location x, y and each snapshot *s_i_* is determined by taking the location of the minimum RIDF (Eq. 3). To weigh the heading direction the ratio *w_si_* was calculated between the minimum RIDF of all snapshots *s_i_* in *S_dmin_* and the current RIDF *d_si_* (Eq. 5). The homing vector *HV⃗* results from the weighted circular mean of the different heading directions *h_si_* (Eq. 6).

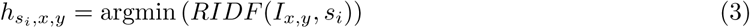

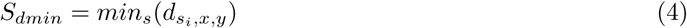

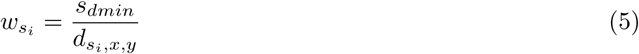

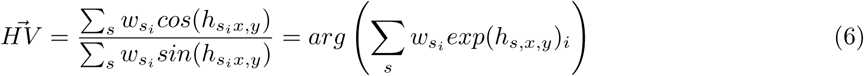

The contrast-weighted-nearness model will use the contrast weighted by the depth of the environment to steer an agent home. The contrast was calculated as the ratio of luminance-amplitude (i.e. standard deviation of the luminance) and luminance-background (i.e. average of the luminance) within a 3×3 pixel window of the snapshot image (i.e. Michelson contrast). As described in [14], we used the rotational similarity function between the current view *I_x,y_* and the memorised view *I_N_* (Eq. 7). As for the brightness-based model, the homing vector (Eq. 6) was computed by the weighted circular means of each vectors derived from each memorised views (Eq. 9).

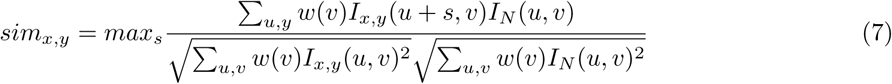

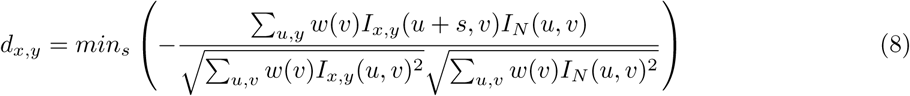

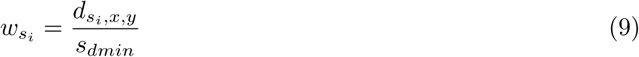

**Table 1.** Statistical results of t-tests

| Group | Condition | Parameter | statistics | p-value | P-value (adjusted) | significance (adjusted) | cohen's d |
| --- | --- | --- | --- | --- | --- | --- | --- |
| $B^+G^+$ | B | Time two locations | 0.698 | 0.491 | 1.474 | ns | -0.179 |
|  | BG | Time two locations | -6.767 | 4.31E-07 | 1.29E-06 | *** | 2.075 |
|  | G | Time two locations | -6.178 | 1.85E-06 | 5.54E-06 | *** | 1.832 |
|  | G and BG | true nest | -0.321 | 0.750 | 2.249 | ns | -0.089 |
|  | G and BG | visual nest | 0.571 | 0.571 | 1.713 | ns | 0.158 |
| $B^-G^+$ | B | Time two locations | 3.589 | 0.001 | 0.004 | ** | -0.910 |
|  | BG | Time two locations | -4.252 | 0.000 | 0.001 | *** | 1.330 |
|  | G | Time two locations | -6.326 | 1.28E-06 | 3.83E-06 | *** | 1.910 |
|  | G and BG | true nest | -0.989 | 0.328 | 0.983 | ns | -0.274 |
|  | G and BG | visual nest | 2.403 | 0.020 | 0.060 | ns | -0.666 |
| Condition | Groups | Parameter | statistics | p-value | P-value (adjusted) | significance (adjusted) | cohen's d |
| BG | $B^-G^+$ and $B^+G^+$ | true nest | 3.324 | 0.002 | 0.005 | ** | 0.922 |
| | $B^-G^+$ and $B^+G^+$ | visual nest | -2.254 | 0.029 | 0.086 | ns | -0.625 |
| G | $B^-G^+$ and $B^+G^+$ | true nest | 1.401 | 0.167 | 0.502 | ns | 0.389 |
| | $B^-G^+$ and $B^+G^+$ | visual nest | 0.899 | 0.373 | 1.119 | ns | 0.249 |

